# Canada’s human footprint reveals large intact areas juxtaposed against areas under immense anthropogenic pressure

**DOI:** 10.1101/2021.06.11.447577

**Authors:** Kristen Hirsh-Pearson, Chris J. Johnson, Richard Schuster, Roger D. Wheate, Oscar Venter

## Abstract

Efforts are underway in Canada to set aside terrestrial lands for conservation, thereby protecting them from anthropogenic pressures. Here we produce the first Canadian human footprint map to identify intact and modified lands and ecosystems. Our results showed strong spatial variation in pressures across the country, with just 18% of Canada experiencing measurable human pressure. However, some ecosystems are experiencing very high pressure, such as the Great Lakes Plains and Prairies national ecological areas which have over 75% and 56% of their areas, respectively, with a high human footprint. In contrast, the Arctic and Northern Mountains have less than 0.02% and 0.2% under high human footprint. A validation of the final map resulted in a Cohen Kappa statistic of 0.911, signifying an ‘almost perfect’ agreement between the human footprint and the validation data set. By increasing the number and accuracy of mapped pressures, our map demonstrates much more widespread pressures in Canada than were indicated by previous global mapping efforts, demonstrating the value in specific national data applications. Ecological areas with immense anthropogenic pressure, highlight challenges that may arise when planning for ecologically representative protected areas.

## Introduction

Global pressures to biodiversity are increasing as human use continues to alter terrestrial ecosystems (Steffen *et al*., 2015; Venter *et al*., 2016a), leading to accelerating biodiversity declines (Maxwell *et al*., 2016; Newbold *et al*., 2015). Anthropogenic pressures to biodiversity are actions taken by humans that have the potential to harm natural systems (Venter *et al*., 2016a). Pressures on a landscape interact with each other in a complex manner and vary in their spatial and temporal scales making their understanding essential for conservation planning (Geldmann *et al*., 2014; Primack, 1993; Tapia-Armijos *et al*., 2017). Identifying the patterns of change in these pressures provides the basis for mitigating environmental damage (Halpern *et al*., 2015; Venter *et al*., 2016a).

When pressures are analysed, especially those from resource development projects, the focus is often on the project in isolation of other developments (Johnson, 2016). By incorporating more than one pressure, it is possible to develop a more complete understanding of the interacting pressures on biodiversity, with the potential to assess the impacts for ecosystem services (Halpern *et al*., 2008). Cumulative pressure mapping allows for the combination of more than one pressure to show the full extent and intensity of anthropogenic pressures (Tapia-Armijos *et al*., 2017)). Cumulative pressure maps, also known as human footprint maps, combine pressures into a single product which can be used for making conservation plans that yield the greatest benefits, including directing development to areas that will cause the least amount of harm (Crain *et al*., 2009; Venter *et al*., 2016a).

As a signatory of the Convention on Biodiversity and its Aichi Biodiversity Targets, Canada’s Target 1 is to protect 17% of terrestrial and 10% of marine areas (MacKinnon *et al*., 2015). With this ambitious conservation target there is a need to better understand the distribution of pressures to natural systems across Canada. A nation-wide map of human pressures is important for identifying the ecosystems that are most intact and the areas with the greatest intensity of human pressures. Intactness is defined as landscapes that maintain biological and ecological function and are mostly free of human disturbances. This definition does not exclude Indigenous peoples and their stewardship practices, but it does exclude large-scale land conversion, human activity and development (Waller and Reo, 2018; Watson *et al*., 2016). Mapping the human footprint will serve as an important step in selecting which areas to protect, restore and sustainably manage.

Canada’s natural systems have a number of pressures that negatively affect biodiversity. Woo-Durand *et al*. (2020) analysed pressures to 820 species identified as “at-risk” by the Committee on the Status of Endangered Wildlife in Canada (COSEWIC). They found that the number of pressures affecting each species has increased significantly from an average of 2.5 to 3.5 between 1999 and 2018. Such findings highlight the need to map pressures in Canada cumulatively and not in isolation (Venter *et al*., 2006; Woo-Durand *et al*., 2020). Nevertheless, no cumulative pressure map covers the entirety of the country. At present, Canada has cumulative pressure maps for parts of the coastal waters (Ban and Alder, 2008; Ban *et al*., 2010; Clarke Murray *et al*., 2015a, 2015b) and two studies covering freshwater (Robb, 2014; Sterling *et al*., 2014). For terrestrial studies, the greatest coverage spans the largest ecological area of Canada, the Boreal/Taiga (Pasher *et al*., 2013). Other terrestrial maps cover sections of western Canada (Mann and Wright, 2018; Shackelford *et al*., 2017), part of eastern Canada and the United States (Woolmer *et al*., 2008) and the whole of Canada to display the number of pollution pressure categories present (McCune *et al*., 2019). In addition to human footprint maps, a map exists showing the presence and absence of access into nature across Canada (Lee and Cheng, 2014). However, a simple map of access does not fully represent the spectrum of human pressures, such as the contrast between large metropolitan cities and smaller resources centric towns (Lee and Cheng, 2014), or landscapes under pressure from resource extraction. The global human footprint map displays Canada as mostly intact (Venter *et al*., 2016a, 2016b), however that includes only a subset of pressures relevant to the country, missing critical data for the Canadian context, such as mining and forestry. Therefore, until a national human footprint is produced, there will continue to be a gap in our understanding for the Canadian human footprint.

Here, we used geospatial techniques to develop a map for Canada that represents nationally-specific pressures that are not incorporated in coarse-scale global maps. Using a higher spatial resolution of 300 metres, we produced the first national terrestrial human footprint of Canada. We visually and quantitatively compared the global and national products and identified improvements and errors in the representation of human pressures. We used high-resolution satellite imagery to validate the accuracy of the final footprint map. As the maintenance of biodiversity and ecosystem services depends on the comprehensive understanding of the full set of overlapping pressures (Sala *et al*., 2000), the results of this project will be important for identifying future conservation lands across Canada as well as ecosystems that are in need of protection and restoration.

## Methods

### Overview

To produce the Canadian human footprint, we adopted the methods originally developed by Sanderson *et al*. (2002) and later refined by Venter *et al*. (2016a, 2016b). The pressures we mapped for Canada were: (1) the extent of built environments; (2) crop land; (3) pasture land; (4) human population density; (5) nighttime lights; (6) railways; (7) roads; (8) navigable waterways; (9) dams and associated reservoirs; (10) mining activity; (11) oil and gas; and (12) forestry. Each anthropogenic pressure was placed on a 0-10 scale to allow for comparison across pressures. Scoring methods were selected from pre-existing peer-reviewed articles following Venter *et al*. (2016a, 2016b) and Woolmer *et al*. (2008) methods and one pressure layer following methods used in Jarvis *et al*. (2010). After scoring, all non-compatible land uses were analysed and adjusted to avoid spatial overlap. Non-compatible land uses included built environments, crop land, mining and pasture land. We eliminated any pixels from the given layers that overlapped with built environments, then did the same operation for crop land and mining. The order of priority, to adjust for spatial overlap, reflected how high up on the 0-10 scale the individual layers placed. To produce the final product of the terrestrial human footprint map of Canada, all the weighted layers were summed together. Individual pressures may overlap spatially and are therefore not mutually exclusive. Thus, each cell could range in value from 0-55 for any given grid cell, representing the observed maximum. The map was generated at a spatial resolution of 300 metres, yielding over 99,000,000 pixels. ArcGIS 10.5.1 and the Lambert Conformal Conic projection were used for all spatial analyses. Specific details on each of the pressure layers are provided in the following sections.

### Built environments

Built environments are lands that are constructed for human activity and include buildings, paved surfaces, and urban areas. Land transformation from built environments leads to habitat loss and fragmentation, changes in nutrient and hydrological flows, reduction of viable habitats for species and decreased temperature regulation and carbon sequestration (Haase, 2009; Tratalos *et al*., 2007).

We acquired data from the 2016 annual crop inventory (Government of Canada; Agriculture and Agri-Food Canada; Science and Technology Branch, 2016), which provides a 30 metre spatial resolution of land-use type and applied the subset of the ‘urban/developed’ lands for the layer. The data does not include Yukon, Northwest and Nunavut territories, and therefore we captured the anthropogenic pressures for the northern territories through other layers such as: population density, nighttime lights and roads. The data are a combination of satellite imagery: Landsat-8, Sentinel-2 and Gaofen-1 for optical imagery with RADARSAT-2 radar imagery, generating an accuracy of at least 85% (Government of Canada; Agriculture and Agri-Food Canada; Science and Technology Branch, 2016). Built environments were assigned a score of 10 (Venter *et al*., 2016a, 2016b).

### Population density

Human population density is linked to biodiversity loss (Cincotta and Engelman, 2000). Presence of high human populations has led to over-hunting, deforestation and introduced species (Prebble and Wilmshurst, 2009). Though Canada generally has a low population density, averaging four people per square kilometre, there have been significant increases in introduced species, over-exploitation and pollution from 1999-2018 (Government of Canada; Statistics Canada, 2017a; Woo-Durand *et al*., 2020).

Human population density was mapped using a subset of the 2016 Canadian Census Data which provides more detailed information than the Gridded Population of the World dataset (Government of Canada; Statistics Canada, 2017a; Venter *et al*., 2016b). The vector layer used was the Census Dissemination Blocks, the smallest unit with an associated population, available through the Geo Suite 2016, a Statistics Canada tool we used for data retrieval (Government of Canada; Statistics Canada, 2016). Following Venter *et al*. (2016a, 2016b), we calculated population density for each block; any block that had more than 1,000 people per square kilometre we assigned a pressure score of 10. For more sparsely populated areas, we logarithmically scaled the pressure score as follows: Pressure Score=3.333∗log(Population density+1)

### Nighttime lights

Nighttime lights captures the sparser electric infrastructure found in rural, suburban and working areas that have an associated pressure on natural environments (Venter *et al*. 2016a, 2016b). The Visible Infrared Imaging Radiometer Suite (VIIRS), mounted on the Sumo National Polar Partnership satellite, provides the means to collect and map low light sources such as nighttime lights (Elvidge *et al*., 2013).

We used an annual composite from 2016 generated by the National Oceanic and Atmospheric Administration (NOAA) to assess nighttime lights. The spatial resolution of the data is 589 metres (15 arc-second geographic grids). For areas above 67N, that were not included in the annual composite, we randomly selected a single date of imagery to fill the northern section and compared it to other dates to make sure it was not an outlier (NOAA, 2019). We then rescaled the data on a 0-10 scale using an equal quintile approach (Venter *et al*. 2016a, 2016b).

### Crop and pasture land

Agriculture is recognised as one of the most important pressures to biodiversity globally (Ricketts and Imhoff, 2003). For the Canadian human footprint, we used the 2016 annual crop inventory which includes pasture, agricultural land, cereals, pulses, oil seeds, vegetables, fruits and other crops (Government of Canada; Agriculture and Agri-Food Canada; Science and Technology Branch, 2016). Satellite imagery from optical (Landsat-8, Sentinel-2 and Geifen-1) and radar (RADARSAT-2) was used to obtain a spatial resolution of 30 metres. There was also ground-truth information provided by several organizations. The provincial accuracy for crop class had a minimum of 86.27% and a maximum accuracy of 94.51% (Government of Canada; Agriculture and Agri-Food Canada; Science and Technology Branch, 2016). we assigned crops a pressure score of 7 (Venter *et al*. 2016a, 2016b).

Pasture lands are areas that are grazed by domesticated livestock. Pastures are often associated with fences, soil compaction, intensive browsing, invasive species and altered fire regimes (Kauffman and Krueger, 1984). Using the annual crop inventory (30 metre spatial resolution) (Government of Canada; Agriculture and Agri-Food Canada; Science and Technology Branch, 2016), We assigned pastures a pressure score of 4 (Venter *et al*. 2016a, 2016b).

### Roads and railways

Roads are linear features that directly convert and fragment habitats. Roads can alter the immediate physical and chemical environments, provide access for human recreation into intact areas, allow for the spread of invasive species and be a sink for populations through vehicle collisions and mortality from construction (Trombulak and Frissell, 2000).

We used the publicly available 2016 National Road Network vector layer produced by Statistics Canada (Government of Canada; Statistics Canada, 2017b). The data are divided into different categories of use: Trans-Canada highway, national highway system, major highway, secondary highways, major streets and all other streets. We adapted the weights developed by Woolmer *et al*. (2008) assessing roads as an access point into intact areas (Table 1).

**Table 1:** Road Pressure Scoring, separated by the different road types to allow for differential scoring. The distances represent the scores associated with each of the buffers.

|  | 0-300m | 300-600m | 600-900m | 900-3000m |
| --- | --- | --- | --- | --- |
| **Road Type** |  |  |  |  |
| Trans-Canada Highway | 10 | 8 | 6 | 4 |
| National Highway and Major Highways | 8 | 6 | 4 | 2 |
| Secondary Highways, Major streets and all other streets | 6 | 4 | 2 | 0 |

Railways provide a direct pressure to the ecosystems that host them, however, in terms of access they differ from roads. For roads and railways, direct pressures exist as a result of the actual footprint such as physical removal of viable habitat or reduction in the quality of it, indirect pressures may present themselves in the form of altering ecological functions, edge effect, reducing connectivity or other human disturbances made possible by the direct pressure (Burton *et al*., 2014). However, discontinued rail lines provide an indirect pressure as they can be used as a means of dispersal of humans and their activities into landscapes. Conversely, operational rails only allow for human access from individual rail stations. We used the publicly available National Railway Network vector layer (Government of Canada; Natural Resources Canada, 2016) and adapted the methods from Woolmer *et al*. (2008) (Table 2).

**Table 2:** Rail Pressure Scoring, separated by operational and discontinued. The distances represent the scores associated with each of the buffers.

|  | 0-300m | 300-600m | 600-900m |
| --- | --- | --- | --- |
| **Rail Type** |  |  |  |
| Operational | 6 | 4 | 0 |
| Discontinued | 6 | 4 | 1 |

### Navigable waterways

Navigable waterways like roads and rails act as means of access to wilderness areas. Canada’s waterways have a long history of human use as they have enabled travel from sea to sea (Brine, 1995). Once the people’s ‘highway’, settlements were formed along the waterways to allow movement and access. Used by First Nations in pre-colonial times, the knowledge was shared when the first European explorers arrived. These waterways were later instrumental in the fur trade (Brine, 1995; O’Donnell, 1989).

We used the dataset generated for navigable coasts for 2009 from the global human footprint with a 1 km^2^ spatial resolution (Venter *et al*., 2016b). The layer included the Great Lakes, as they can act like inland seas and was generated using distance to settlements, stream depth and hydrological data (Venter *et al*., 2016b). We found the centreline of the waterway then weighted them to follow the other access-based layers (Table 3).

**Table 3:** Navigable Waterway Pressure Scoring. The distances represent the scores associated with each of the buffers.

|  | 0-300m | 300-600m | 600-900m |
| --- | --- | --- | --- |
| Navigable Waterways | 6 | 4 | 2 |

### Dams and reservoirs

Dams directly change hydrology of the areas and they modify the environment, often producing human-made flooded reservoirs (Woolmer *et al*., 2008). The vector dataset was obtained from ‘Large Dams and Reservoirs of Canada’ (Global Forest Watch Canada, 2010). We mapped the dam itself just as we would a built environment, scoring it as 10 (Venter *et al*., 2016a, 2016b; Woolmer *et al*., 2008). We scored dams and associated reservoirs in the same manner as navigable waterways given that they can provide additional access to areas by watercraft (Table 3).

### Mining

Mining often alters topography, watercourses and removes topsoil as a form of land conversion. Mining can be a point source for air and water pollution (Woolmer *et al*., 2008). We used the mines and minerals dataset, updated in 2015, to obtain all active mines in Canada. The data were discrete points in vector format (Government of Canada; Natural Resources Canada, 2017). We placed the mineral groups in their designated categories: open large, open small, underground large and underground small (WWF Canada, 2003). For the minerals that were not previously classified by Woolmer *et al*. (2008) we consulted with an expert to determine if the mineral group would be mined underground or in an open pit (McGill, 2018). Once confirmed to be open or underground, we placed them all in the small category, for open pit and underground mining, as a way to make sure we did not over-estimate the pressure. The scoring from Woolmer *et al*. (2008) was used for mines (Table 4).

**Table 4:**
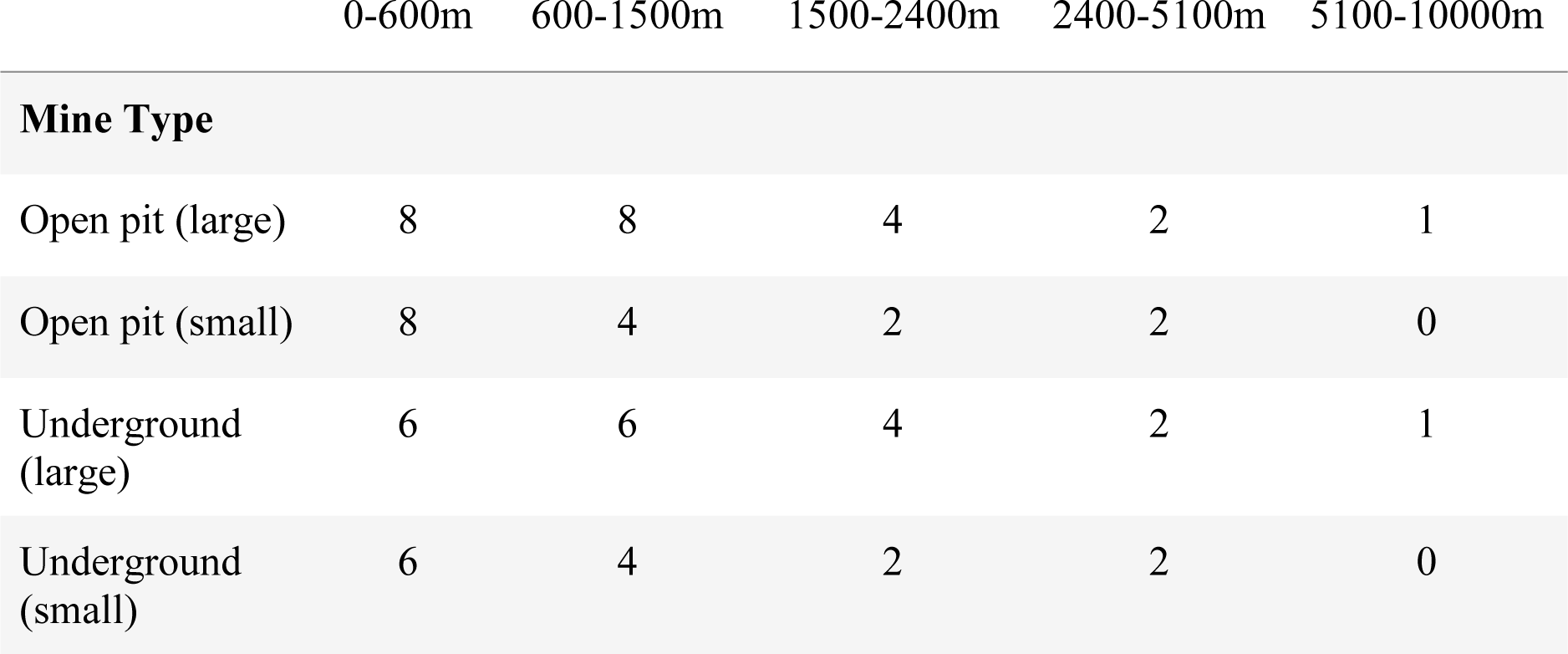
Mines Pressure Scoring, separated by the designated mining categories. The distances represent the scores associated with each of the buffers.

### Oil and gas

Oil and gas production have a number of associated pressures to nature such as wildlife mortality, habitat fragmentation and loss, noise and light pollution, introduction of invasive species and sedimentation of waterways (Brittingham *et al*., 2014; Jones *et al*., 2015). The mines and minerals dataset, updated in 2015, was used as it lists active oil and gas fields. The data were discrete points in vector format (Government of Canada; Natural Resources Canada, 2017). The direct pressures from oil and gas have been found to be highly localized, therefore, we adapted the scoring method using a 10 to 0 scale to score the linear circular decay out to five kilometres away from the site centre (Jarvis *et al*., 2010).

### Forestry

Forestry operations alter the forest structure by changing stand dynamics and age (Freedman *et al*., 1994). Clear cut forestry can remove habitat for species dependent on old trees, deadwood and tree cavities and, by altering paths of travel and allowing for deep snow to form. Forestry operations could also introduce species and allow for more access for recreation including hunting through the creation of forestry roads (Freedman *et al*., 1994).

The forest-harvest data were obtained from an annual forest disturbance characterization project for Canada that has a 30-metre spatial resolution (White *et al*., 2017). The timescale of the harvest recorded was from 1985-2015. We separated fresh clear cuts and areas that have reached their free-to-grow state, as they offer different habitat qualities (Bergeron *et al*., 2011). We selected 12 years as a common value for free-to-grow, so anything from 0-12 years would be considered early regenerating forest (Lieffers *et al*., 2002; Smith, 1983). We adapted the scoring from Woolmer *et al*. (2008) with early regeneration scored as 4 and older regeneration as 2 (Woolmer *et al*., 2008).

### Technical Validation

Following the methods used by Venter *et al*. (2016a, 2016b), a single person used high resolution satellite imagery to visually identify human pressures within 5,000, 1-km^2^ randomly located sample plots. Using World Imagery, available through ArcGIS, the 5,000 plots had a median resolution of 0.5 metres and a median acquisition year of 2014 (ArcGIS, n.d.).

We used Venter *et al*. (2016a, 2016b) methods to develop a standardized key to visually interpret the pressures. For the eight pressures that both our Canadian human footprint and the global human footprint had in common we mimicked their scoring, but for the new pressures included in our study we simply followed their standards for linear or polygons features (Supplementary Information, S1). Interpretations were marked if they were ‘certain’ or ‘uncertain’; in our case 254 plots were ‘uncertain’ and therefore discarded, leaving 4,746 validation plots. Generally, plots were classified as ‘uncertain’ for two main reasons: due to inadequate resolution of the imagery (15 metres) so it was not clear if there were any pressures present on the land, or because of cloud cover obscuring some or all of the image. The plots that were retained for the visual scoring were all ‘certain’ and we therefore consider the in-situ pressures for the plot as true. The mean human footprint score for the 1-km^2^ plots were determined in ArcGIS, then both the visual and human footprint scores were normalized on a 0-1 scale.

The root mean squared error (Chai and Draxler, 2014) and the Cohen kappa statistic of agreement (Viera and Garrett, 2005) were used to quantify the level of agreement between the Canadian footprint map and the validation dataset. The root mean squared error measures the differences between the values calculated in the human footprint and the visual scores from the validation. As the error is squared, outliers are emphasized with this statistical calculation. The Cohen kappa statistic of agreement expresses the agreement between the human footprint scores and the visual interpretation scores considering the potential that agreement or disagreement may occur by chance. Following previous analyses (Venter *et al*., 2016a, 2016b), visual plots that were within 20% of the human footprint plots scores were considered a match for the Cohen kappa statistic.

## Results

### 1. The Canadian human footprint

Canada has an area-weighted average human footprint score of 1.48, and the maximum observed score for the country is 55 out of a theoretical 66. The pressures across Canada display strong spatial patterns, showing higher values in Southern Canada where the majority of the country’s population lives (Fig. 1). With the 12 pressures included, we found that 82% of Canada’s land areas had a human footprint score of less than 1, and therefore were considered intact (Allan *et al*., 2017). In this context, intact is defined as landscapes that are mostly free of the 12 human disturbances we mapped. To conceptualize this definition of intact, cells that had a population density of one or more people per square kilometre obtained a pressure score of one or above and were therefore not considered intact. However, pressures such as seismic lines, pollution or invasive species were not mapped and may be present in areas that we identified as intact. The low human footprint state was defined as areas where the human footprint score was between one and four. The upper limit was determined based on the assignment of a score of four for pasture land, which would often have fences fragmenting the connectedness (Venter *et al*., 2016a). Approximately 5% of Canada was classified in the low human footprint state. The moderate human footprint areas had scores between four and 10 and covered 7% of the country. The areas of high human footprint, with a value of 10 or higher, covered 6% of Canada and highlighted areas with multiple overlapping pressures to biodiversity (Fig. 1).

**Figure 1:**
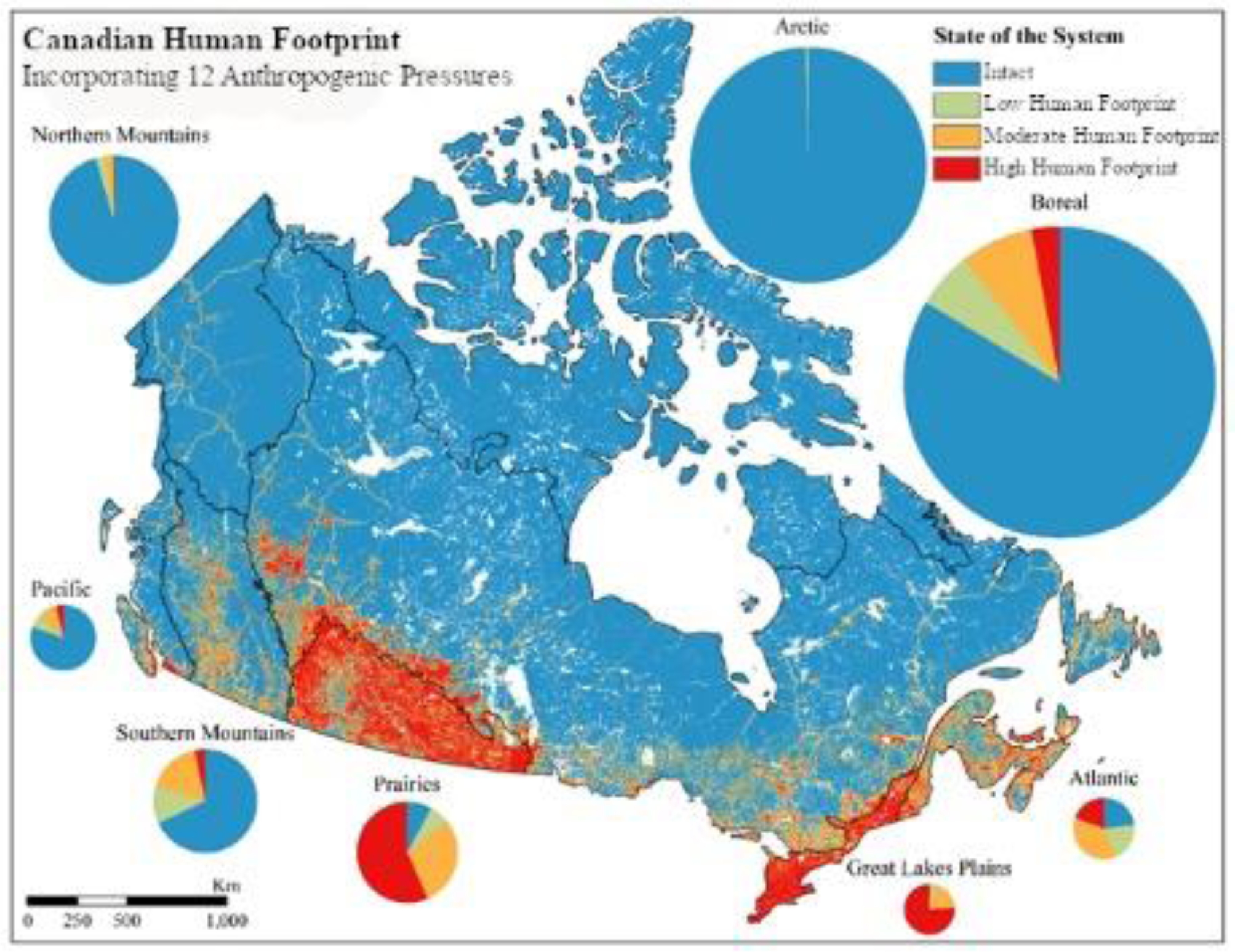
Human footprint map of Canada showing the state of the system for COSEWIC national ecological areas. Pie chart sizes represent the approximate proportions each ecological area covers of Canada. The footprint represents 12 anthropogenic pressures: built environments, population density, nighttime lights, crop land, pasture land, roads, railways, navigable waterways, dams and associated reservoirs, mines, forestry and oil and gas.

We used national ecological areas defined by COSEWIC as a means of comparing the different ecological regions of Canada (COSEWIC, 2018). The human footprint differs markedly across those areas, with 84% of the Boreal ecological area, which covers the largest extent of Canada, still being intact. The Great Lakes Plains, the smallest ecological area, has 76% in the high human footprint category, being the largest percentage in the high category compared to all other ecological areas. The Prairies follow the Great Lakes Plains as the second largest values in the high human footprint category with 57%. The Great Lakes Plains has the smallest percentage in the intact category with a value of 0.6% followed by the Prairies with 8%. Conversely, the Arctic, which is the second largest ecological area, is over 99% intact.

The pressure layer that contributes the most towards the mean human footprint of Canada is roads with a mean human footprint score of 0.72 (Fig. 2) and covering over 1,000,000 km^2^. Crop land is the second most prevalent pressure with a mean human footprint score of 0.27. The only other pressure above 0.10 was population density with a value of 0.20. In terms of extent, population density covers just under one third of Canada, an area of 3,200,000 km^2^. While nighttime lights cover over 200,000 km^2^, they have a relatively small mean human footprint of 0.01.

**Figure 2:**
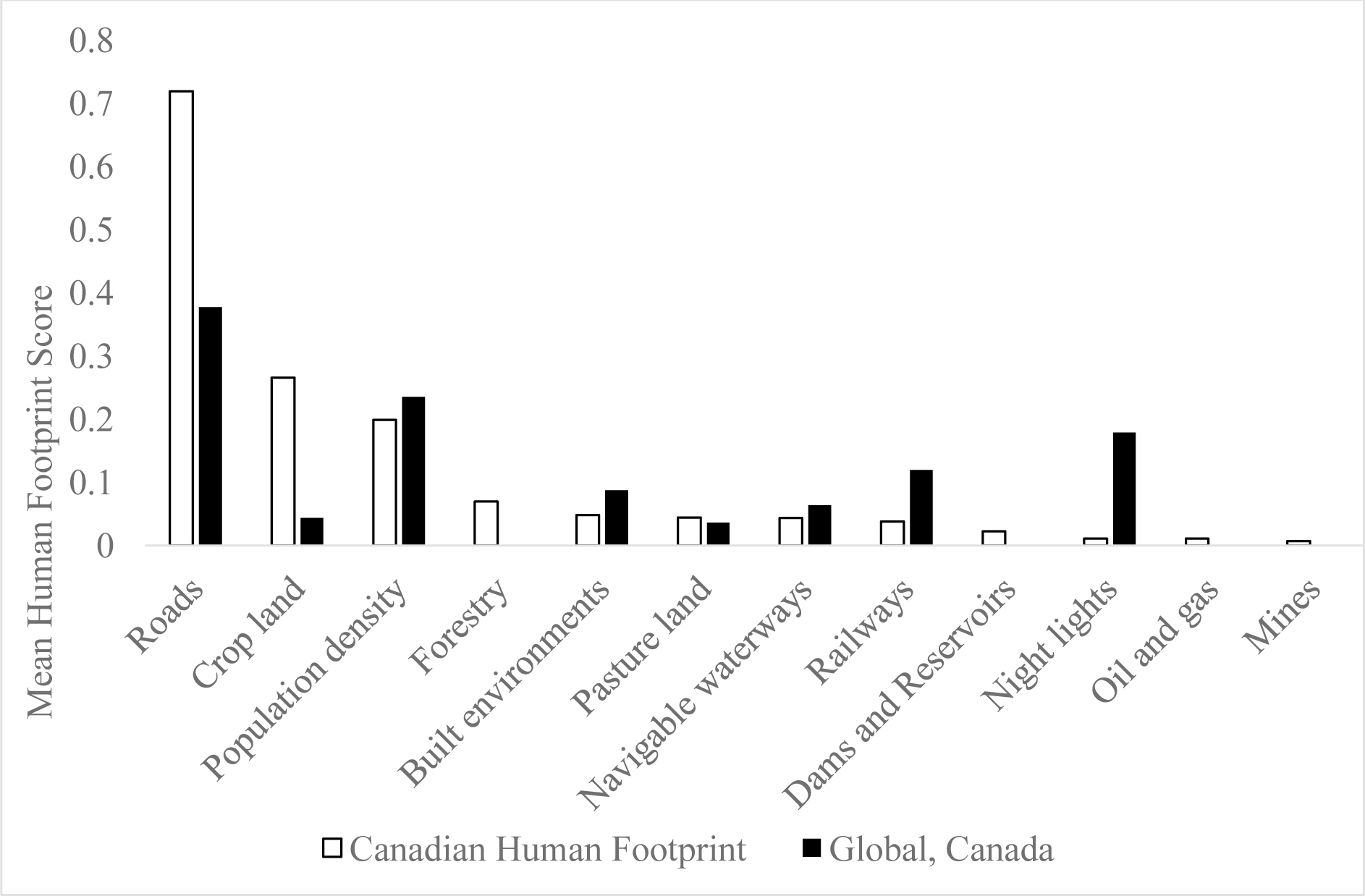
Mean human footprint scores for each of the pressures included in the Canadian human footprint and the Global Human Footprint. ‘Canadian human footprint’ (white) are from the results produced in this project, the ‘Global, Canada’ (black) is from the global human footprint product clipped to Canada for comparison.

### 2. The Canadian versus the Global human footprint

Visually comparing the global human footprint (Venter *et al*., 2016a) to the national version at a broad scale shows similarities in the spatial patterns of anthropogenic pressures (Supplementary Information, S2). Closer examination shows a number of variations in the details. In agricultural areas, such as the prairies ecological region, the Canadian human footprint shows a higher concentration of pressures than the global one (Fig. 3 a, b, c). For urban areas, the Canadian human footprint captures the distinction between areas such as parks, urban areas and industrial areas showing a lower human footprint score than the global one (Fig. 3 d, e, f). In natural resource intensive areas, higher scores for the Canadian human footprint are present compared to the global product that missed these features across Canada. For example, in the boreal ecological area, forestry harvest and infrastructure from oil and gas could be included with the Canadian human footprint (Fig. 3 g, h, i).

**Figure 3:**
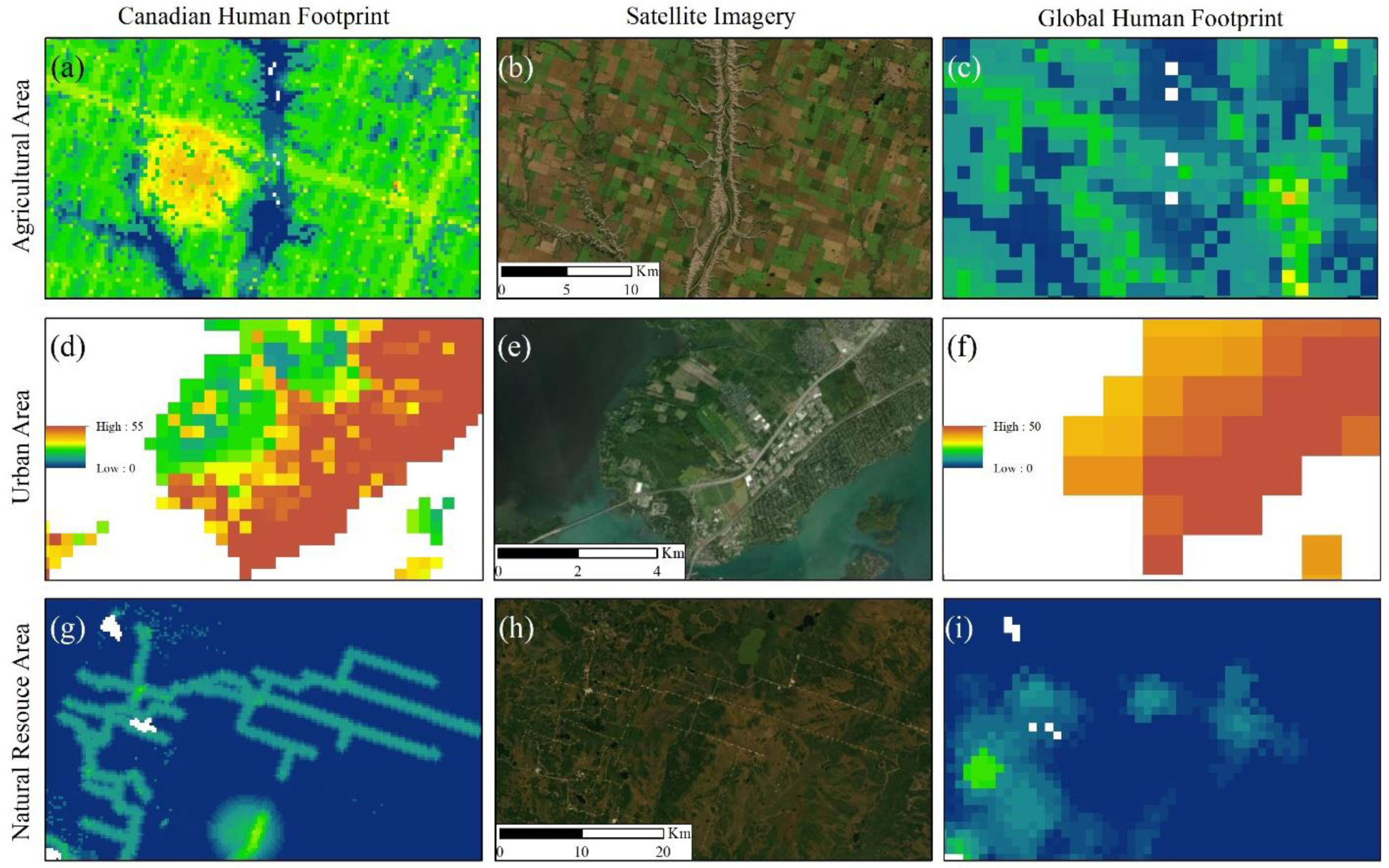
Visual comparison between the Canadian human footprint (first column) the high resolution satellite imagery (second column) and the Global human footprint (third column). The first row, Agricultural Area (a, b, c), is located in the prairies ecological area. The second row, Urban Area (d, e, f), shows the western part of the island of Montreal which is located in the Great Lakes Plains ecological area. The third row, Natural Resource Area (g, h, i), located in the Boreal ecological area, looks at a natural resource intensive area where forestry cutblocks and oil and gas infrastructure are present. The legend for column one is found in pane ‘d’ and for column three in pane ‘f’. The scale bar for each row is found in the second column. The source for first column is from this project, second column is from the high resolution imagery basemap option in ArcGIS and the third column from Venter *et al*. (2016a, 2016b).

When mapping nationally explicit data the greatest improvements to the global datasets were found with the National Roads Network and the Annual Crop Inventory. The global human footprint scores roads within Canada as 50% less of a pressure than the Canadian human footprint. The Annual Crop Inventory that was used for mapping crop land for Canada captured over 285,000 km^2^ more than the global product (Fig. 2).

### 3. Validation results

Our validation shows a strong agreement between the Canadian human footprint measure of pressures and the pressures scored using visual interpretation of high-resolution images. The root mean squared error for 4,746 validation 1-km^2^ plots was 0.07 on a normalized 0–1 scale (Chai and Draxler, 2014). The Cohen Kappa statistic was 0.911, signifying ‘almost perfect’ agreement between the human footprint and the validation data set (Landis and Koch, 1977; Viera and Garrett, 2005). We scored 40 of the validation plots as having a pressure score 20% higher than the initial visual interpretation (false positive) and 113 20% lower (false negative). The remaining 4,593 plots (96.8%) were within 20% agreement. While the results from the validation represent almost perfect agreement, it appears from the higher false-negative rate that the human footprint map may be underrepresenting the pressure scores across some proportion of the country. The maps should therefore be considered as conservative estimates of anthropogenic pressures on the environment (Fig. 4). When applying a more rigorous threshold for agreement, within 15% of one another, we found that the Cohen Kappa statistic was of substantial strength with a score of 0.772. When applying a less rigorous threshold of 25%, the Cohen Kappa statistic increased to 0.952 (almost perfect strength) (Supplementary Information, S3).

**Figure 4:**
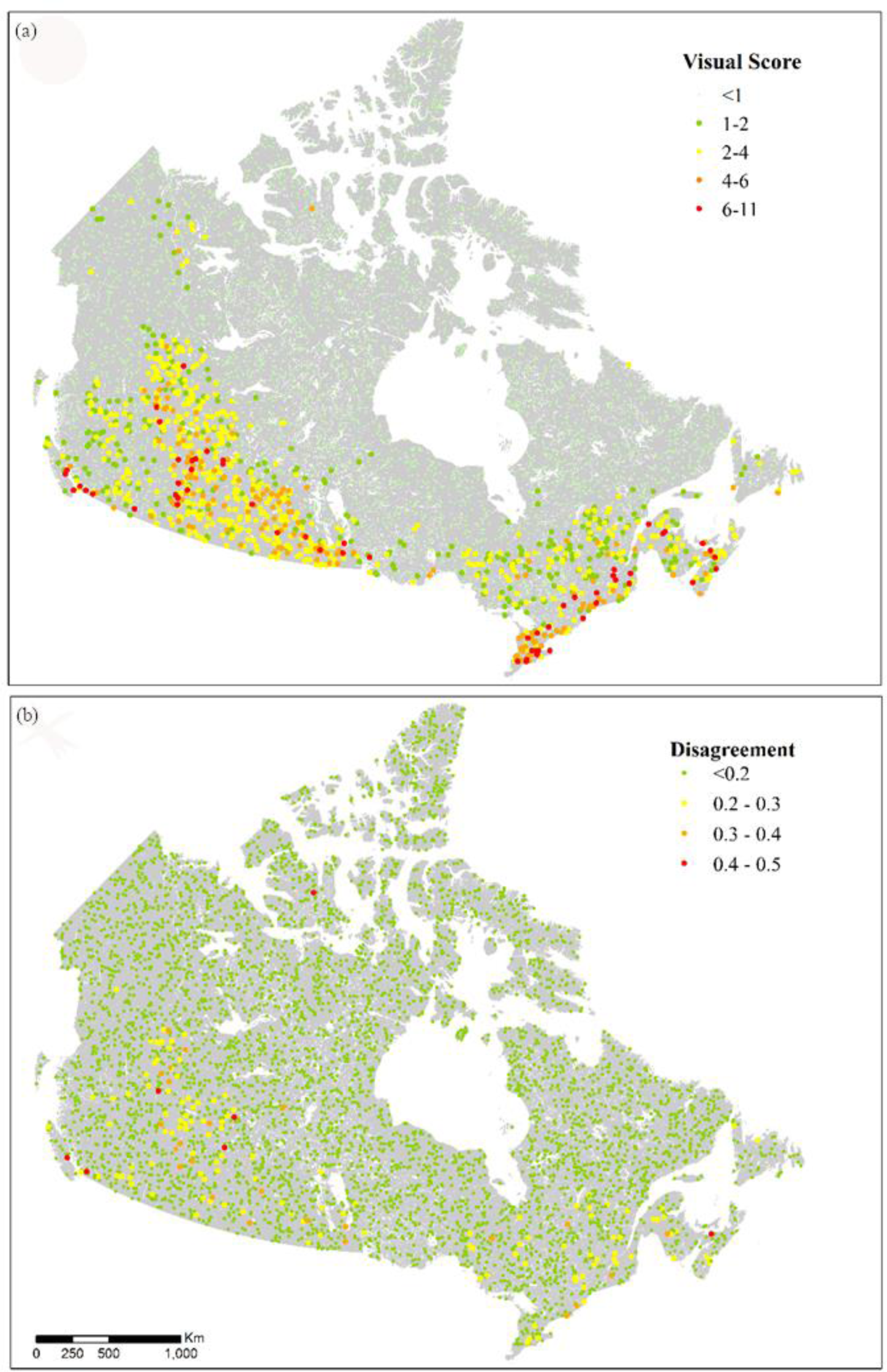
Results from the 4,746 x 1km^2^ validation plots interpreted and scored following Supplementary Information, S1. (a) the visual interpretation score assigned and location for plots, and (b) the disagreement between the Canadian human footprint score and the visual interpreted score for validation normalized on a 0-1 scale.

We compared the validation results for the Canadian human footprint with those of the global human footprint clipped to Canada. The global human footprint obtained a root mean squared error of 0.10 on a normalized 0–1 scale for the same validation plots (Chai and Draxler, 2014). For the Cohen Kappa statistic, the value was 0.762 using 20% agreement, which is considered substantial agreement between the human footprint and the validation data set, demonstrating lower agreement than the Canadian product (Landis and Koch, 1977; Viera and Garrett, 2005).

## Discussion

This is the first undertaking to produce a continuous measure of human pressures across Canada, which we term the Canadian human footprint. While we find that the large majority of Canada is still considered intact (82%), by our definition, some ecosystems are still exposed to numerous and intense pressures. Our dataset improves upon the global product by increasing the number of relevant pressures measured and by rescaling to a finer resolution using national datasets. Understanding where there are overlapping pressures on the natural system provides more insight for preventing and mitigating pressures to biodiversity than an access map of Canada that acts as a binary presence or absence of access.

### Intactness in Canada

Intact areas worldwide are crucial for conserving threatened biodiversity (Di Marco *et al*., 2019), yet they experience increasing pressures from human land use. There is therefore a need to protect intact areas to help conserve biodiversity and ecosystem services. Furthermore, the importance of large-scale and intact ecosystems is increasing as these areas become rarer (Watson *et al*., 2016). When applying the 12 anthropogenic pressures, to the eight national ecological areas in Canada, we find that five of those areas have a terrestrial land mass that is over 50% in an intact state. In particular, the Arctic has over 99%, the Northern Mountains has over 95% and the Boreal follows with over 83%. This demonstrates promising opportunities for the three largest ecological areas in Canada. However, the Boreal is experiencing significant forest loss and degradation from natural resource exploration, industrial forestry, rapid climate change and anthropogenic fires (Watson *et al*., 2016). Although faced with criticism, the Canadian Boreal Forest Conservation Framework provides an outline on how to protect at least 50% of the forest through a network of connected protected areas, to prevent excessive degradation, which is crucial to protect wilderness areas of Canada (Boreal Leadership Council, 2003; Nishnawbe Aski Nation, n.d.).

Canada has little intact areas left in three of the eight ecological areas (Great Lakes Plains, Prairies and Atlantic). Our human footprint shows where species are experiencing the most anthropogenic pressures and would likely have the least intact natural ecosystem for disturbance sensitive species. However, it is known that certain species can thrive in large cities and built environments (Sanderson *et al*., 2002). As mentioned above, Canada’s Target 1 of 17% conservation of terrestrial lands requires those areas to be representative of the county’s ecosystems (Convention on Biological Diversity, 2020). With the Atlantic, Prairies and Great Lakes Plains areas containing less than 24%, 8% and 1% of intact lands respectively, it is unclear how Canada will develop protected areas that represent the ecosystems in those regions, without restoration.

### Pressures on Biodiversity

One of the most prevalent pressures to intactness and biodiversity are roads. The existence and expansion of roads to connect communities and resource areas are direct and indirect pressures to ecosystems, such as fragmenting habitats and by providing a means of access into intact areas (Lee and Cheng, 2014; Sanderson *et al*., 2002). For conservation efforts, roads are one of the important pressures to address (van der Marel *et al*., 2020), especially in Canada where we find roads are the most prevalent pressure.

Conservation planning recognises the need to understand the patterns of pressures and how they interact (Margules and Pressey, 2000; Tulloch *et al*., 2015). When assessing national at-risk species, the Canadian Living Planet Index found a 59 per cent decline in these at-risk populations between 1970 and 2016 (WWF Canada, 2020). A global analysis of over 8,000 threatened or near-threatened species found that overexploitation was the most important pressure, followed by agricultural activities and then urban development (Maxwell *et al*., 2016). For our analysis, those pressures were represented by the footprint of crop land, forestry, built environments and pasture land, all of which fall within the top six mean human footprint scores for Canada. With Canada’s “at-risk” species facing more than one pressure (Woo-Durand *et al*., 2020), the utility of the Canadian human footprint, which includes the pressures that are most affecting biodiversity (Maxwell *et al*., 2016), is an important tool for conservation planning and the mitigation of such pressures.

### Data comparison: Global vs. Canadian

While the overall intensity, or the mean human footprint score, remained low for Canada, we found several differences when comparing global and national human footprints. The most significant difference in the mean human footprint score was found in nighttime lights. Nighttime lights for Canada had a mean human footprint score 18 times less than in the global human footprint. The reduction in score from the global to the national product is the result of using more recent and higher resolution imagery that addresses saturation and spillage observed with the global product (Elvidge *et al*., 2013).

Producing the human footprint for Canada allows us to include datasets that are nationally relevant and offer more information and detail than many of the global footprint maps. The largest increase in mean human footprint score comes from crop land which has a mean human footprint score over six times more extensive than that in the global product. The improved accuracy for mapping crop land could be part of the reason we see higher footprint values in the Prairies when compared to the global product, as the Prairies are a large agricultural centre for the country. Furthermore, the Canadian dataset for roads allowed for the inclusion of minor roads which the global dataset could not include (Venter *et al*., 2016b). The Canadian data also led to a near doubling of the mean human footprint score for roads when compared to Canada’s score with the global data. However, there is still room for improvement in mapping linear infrastructure in Canada. When we compare the national roads with some provincial road data, we find that the national data do not capture all the resource roads and some of the smaller roads that are mapped at a provincial or territorial scale.

The global human footprint and the Canadian human footprint show the same overall patterns of pressures. However, we find more disagreements in areas where there are more cumulative pressures. By developing a finer resolution national product with Canadian data, we can measure the improvements from global human footprints and confirm the soundness of our human footprint with the almost perfect validation score. This demonstrates the importance of national studies for conservation of biodiversity and ecosystem services (Woolmer *et al*., 2008).

### Future directions

This is the first national product for the Canadian human footprint, with room for future refinements. Firstly, linear features besides roads, such as seismic lines or outdoor recreation such as trails, should be included if possible, in future revisions. These features appeared in approximately 1% of the validation plots but were not mapped as there were no national datasets for oil and gas exploration and recreation. Also, recreation more broadly can have significant impacts (Mullins and Wright, 2016), and should be included as data become available. These data should be a priority for future improvements to our work. Other pressures such as extreme weather and introduced species are important, but inherently difficult to map (Venter *et al*., 2006; Woo-Durand *et al*., 2020).

Secondly, the built environments dataset did not cover the full extent of the country (only to 59 degrees north), therefore the theoretical max north of 59 degrees latitude is 10 lower for the human footprint, leading to a potential underestimation of anthropogenic pressure. This is unlikely to be a major omission, as these pressures are sparse or absent above this latitude. Despite lower population density in the north, natural resource exploration has increased, bringing with it more temporary workers and work camps (Ensign *et al*., 2014). Thirdly, the datasets we used to map mining and oil and gas only provided point features. Further efforts are needed to develop complete polygon boundary and associated linear features that more accurately represent the geographic extent of the pressure. Lastly, our product is not immune to the limitations of spatial analyses such as mixed pixel problems, that arise when resampling to the resolution of the project; and the assumption of linear and consistent responses of ecosystems to pressures (Halpern and Fujita, 2013). While there is certainly scope for further refinement, we do note that the validation of Canadian human footprint revealed in a much closer agreement between our dataset and actual observable pressures than previous efforts in other jurisdictions (Kennedy *et al*., 2019; Venter *et al*., 2016a; Williams *et al*., 2020).

## Conclusion

Our Canadian human footprint map provides a baseline from which we can measure changes in human pressures across the country and into the future. Such information is critical for assessing the effectiveness of national and international policies and agreements designed to maintain biodiversity and expand Canada’s conservation and protected lands. Our cumulative pressure map provides the first step towards being able to translate mapped pressures to the impacts of those pressures for biodiversity and ecosystem services. We demonstrate that Canada does contain large intact areas in line with Watson *et al*. (2016) who identified North America as with a critical stronghold for large tracts of intact wilderness. Understanding how Canada’s intact lands are lost through cumulative human activities and associated pressures is crucial for the future prevention and mitigation of biodiversity loss and degradation of ecosystem services in a country where large intact areas still remain.

## Acknowledgements

Project funding was provided by University of Northern British Columbia and Natural Science and Engineering Research Council (NSERC). We are appreciative to M. Hirsh-Pearson, D. Pinkham, L. Spooner, the Conservation Solutions Lab members for the helpful comments during the production of the manuscript. We thank S. Emmons and P. Bai for GIS support and troubleshooting.

## References

Allan, J.R., Venter, O., Watson, J.E.M., 2017. Temporally inter-comparable maps of terrestrial wilderness and the Last of the Wild. Sci. Data 4, 170187. https://doi.org/10.1038/sdata.2017.187

ArcGIS, n.d. World_Imagery (MapServer). ESRI, Redlands, CA [WWW Document]. URL http://services.arcgisonline.com/ArcGIS/rest/services/World_Imagery/MapServer (accessed 3.6.20).

Ban, N., Alder, J., 2008. How wild is the ocean? Assessing the intensity of anthropogenic marine activities in British Columbia, Canada. Aquat. Conserv. Mar. Freshw. Ecosyst. 18, 55– 85. https://doi.org/10.1002/aqc.816

Ban, N.C., Alidina, H.M., Ardron, J.A., 2010. Cumulative impact mapping: Advances, relevance and limitations to marine management and conservation, using Canada’s Pacific waters as a case study. Mar. Policy 34, 876–886. https://doi.org/10.1016/j.marpol.2010.01.010

Bergeron, D.H., Pekins, P.J., Jones, H.F., Leak, W.B., 2011. Moose browsing and forest regeneration: A case study in Northern New Hampshire. Alces J. Devoted Biol. Manag. Moose 47, 39–51.

Boreal Leadership Council, 2003. Canadian Boreal Forest Conservation Framework 8.

Brine, R.H., 1995. Canada’s forgotten highway. Whaler Bay Press, Galiano, B.C.

Brittingham, M.C., Maloney, K.O., Farag, A.M., Harper, D.D., Bowen, Z.H., 2014. Ecological Risks of Shale Oil and Gas Development to Wildlife, Aquatic Resources and their Habitats. Environ. Sci. Technol. 48, 11034–11047. https://doi.org/10.1021/es5020482

Burton, A.C., Huggard, D., Bayne, E., Schieck, J., Sólymos, P., Muhly, T., Farr, D., Boutin, S., 2014. A framework for adaptive monitoring of the cumulative effects of human footprint on biodiversity. Environ. Monit. Assess. 186, 3605–3617. https://doi.org/10.1007/s10661-014-3643-7

Chai, T., Draxler, R.R., 2014. Root mean square error (RMSE) or mean absolute error (MAE)? - Arguments against avoiding RMSE in the literature. Geosci. Model Dev. 7, 1247.

Cincotta, R.P., Engelman, R., 2000. Nature’s Place: Human Population and the Future of Biological Diversity. Population Action International, Washington, DC.

Clarke Murray, C., Agbayani, S., Alidina, H.M., Ban, N.C., 2015a. Advancing marine cumulative effects mapping: An update in Canada’s Pacific waters. Mar. Policy 58, 71– 77. https://doi.org/10.1016/j.marpol.2015.04.003

Clarke Murray, C., Agbayani, S., Ban, N.C., 2015b. Cumulative effects of planned industrial development and climate change on marine ecosystems. Glob. Ecol. Conserv. 4, 110– 116. https://doi.org/10.1016/j.gecco.2015.06.003

Convention on Biological Diversity, 2020. The Convention on Biological Diversity. Secretariat of the Convention on Biological Diversity, Montreal, Qc [WWW Document]. URL https://www.cbd.int/convention/ (accessed 7.2.20).

COSEWIC, 2018. Cosewic / Cosepac - Guidelines for Recognizing Designatable Units. COSEWIC, Gatineau, QC [WWW Document]. URL http://www.cosewic.ca/index.php/en-ca/reports/preparing-status-reports/guidelines-recognizing-designatable-units (accessed 6.26.20).

Crain, C.M., Halpern, B.S., Beck, M.W., Kappel, C.V., 2009. Understanding and Managing Human Threats to the Coastal Marine Environment. Ann. N. Y. Acad. Sci. 1162, 39–62. https://doi.org/10.1111/j.1749-6632.2009.04496.x

Di Marco, M., Ferrier, S., Harwood, T.D., Hoskins, A.J., Watson, J.E.M., 2019. Wilderness areas halve the extinction risk of terrestrial biodiversity. Nature 573, 582–585. https://doi.org/10.1038/s41586-019-1567-7

Elvidge, C.D., Baugh, K.E., Zhizhin, M., Hsu, F.-C., 2013. Why VIIRS data are superior to DMSP for mapping nighttime lights. Proc. Asia-Pac. Adv. Netw. 35, 62. https://doi.org/10.7125/APAN.35.7

Ensign, P.C., Giles, A., Oncescu, J., 2014. Natural Resource Exploration and Extraction in Northern Canada: Intersections with Community Cohesion and Social Welfare. J. Rural Community Dev. 9, 112–133.

Freedman, B., Woodley, S., Loo, J., 1994. Forestry practices and biodiversity, with particular reference to the Maritime Provinces of eastern Canada. Environ. Rev. 2, 33–77. https://doi.org/10.1139/a94-003

Geldmann, J., Joppa, L.N., Burgess, N.D., 2014. Mapping Change in Human Pressure Globally on Land and within Protected Areas. Conserv. Biol. 28, 1604–1616. https://doi.org/10.1111/cobi.12332

Global Forest Watch Canada, 2010. Large Dams and Reservoirs of Canada.

Government of Canada; Agriculture and Agri-Food Canada; Science and Technology Branch, 2016. Annual Crop Inventory.

Government of Canada; Natural Resources Canada, 2017. Principal Mineral Areas, Producing Mines, and Oil and Gas Fields in Canada.

Government of Canada; Natural Resources Canada, 2016. National Railway Network. Government of Canada; Statistics Canada, 2017a. Population and Dwelling Count Highlight Tables, 2016 Census. Government of Canada, Ottawa, On [WWW Document]. URL https://www12.statcan.gc.ca/census-recensement/2016/dp-pd/hlt-fst/pd-pl/Table.cfm?Lang=Eng&T=101&S=50&O=A (accessed 5.9.19).

Government of Canada; Statistics Canada, 2017b. Road Network File 2016.

Government of Canada; Statistics Canada, 2016. Geosuite, Government of Canada, Ottawa, On [WWW Document]. URL https://geosuite.statcan.gc.ca/geosuite/en/index (accessed 6.25.20).

Haase, D., 2009. Effects of urbanisation on the water balance – A long-term trajectory. Environ. Impact Assess. Rev. 29, 211–219. https://doi.org/10.1016/j.eiar.2009.01.002

Halpern, B.S., Frazier, M., Potapenko, J., Casey, K.S., Koenig, K., Longo, C., Lowndes, J.S., Rockwood, R.C., Selig, E.R., Selkoe, K.A., Walbridge, S., 2015. Spatial and temporal changes in cumulative human impacts on the world’s ocean. Nat. Commun. 6, 7615. https://doi.org/10.1038/ncomms8615

Halpern, B.S., Fujita, R., 2013. Assumptions, challenges, and future directions in cumulative impact analysis. Ecosphere 4, 1–11. https://doi.org/10.1890/ES13-00181.1

Halpern, B.S., McLeod, K.L., Rosenberg, A.A., Crowder, L.B., 2008. Managing for cumulative impacts in ecosystem-based management through ocean zoning. Ocean Coast. Manag. 51, 203–211. https://doi.org/10.1016/j.ocecoaman.2007.08.002

Jarvis, A., Touval, J.L., Schmitz, M.C., Sotomayor, L., Hyman, G.G., 2010. Assessment of threats to ecosystems in South America. J. Nat. Conserv. 18, 180–188. https://doi.org/10.1016/j.jnc.2009.08.003

Johnson, C.J., 2016. Defining and Identifying Cumulative Environmental, Health, and Community Impacts, in: The Integration Imperative - Cumulative Environmental, Community and Health Effects of Multiple Natural Resource Developments. Springer International Publishing, pp. 21–45.

Jones, N.F., Pejchar, L., Kiesecker, J.M., 2015. The Energy Footprint: How Oil, Natural Gas, and Wind Energy Affect Land for Biodiversity and the Flow of Ecosystem Services. BioScience 65, 290–301. https://doi.org/10.1093/biosci/biu224

Kauffman, J.B., Krueger, W.C., 1984. Livestock Impacts on Riparian Ecosystems and Streamside Management Implications…A Review. Rangel. Ecol. Manag. J. Range Manag. Arch. 37, 430–438.

Kennedy, C.M., Oakleaf, J.R., Theobald, D.M., Baruch-Mordo, S., Kiesecker, J., 2019. Managing the middle: A shift in conservation priorities based on the global human modification gradient. Glob. Change Biol. 25, 811–826. https://doi.org/10.1111/gcb.14549

Landis, J.R., Koch, G.G., 1977. The Measurement of Observer Agreement for Categorical Data. Biometrics 33, 159–174. https://doi.org/10.2307/2529310

Lee, P., Cheng, R., 2014. Human Access in Canada’s Landscape, Global Forest Watch Canada Bulletin. Global Forest Watch Canada.

Lieffers, V.J., Pinno, B.D., Stadt, K.J., 2002. Light dynamics and free-to-grow standards in aspen-dominated mixedwood forests. For. Chron. 78, 137–145. https://doi.org/10.5558/tfc78137-1

MacKinnon, D., Lemieux, C.J., Beazley, K., Woodley, S., Helie, R., Perron, J., Elliott, J., Haas, C., Langlois, J., Lazaruk, H., Beechey, T., Gray, P., 2015. Canada and Aichi Biodiversity Target 11: understanding ‘other effective area-based conservation measures’ in the context of the broader target. Biodivers. Conserv. 24, 3559–3581. https://doi.org/10.1007/s10531-015-1018-1

Mann, J., Wright, P., 2018. The human footprint in the Peace River Break, British Columbia (No. 2), Technical Report Series. Natural Resources and Environmental Studies Institute, University of Northern British Columbia, Prince George, BC.

Margules, C.R., Pressey, R.L., 2000. Systematic conservation planning. Nature 405, 243–253. https://doi.org/10.1038/35012251

Maxwell, S.L., Fuller, R.A., Brooks, T.M., Watson, J.E.M., 2016. Biodiversity: The ravages of guns, nets and bulldozers. Nat. News 536, 143. https://doi.org/10.1038/536143a

McCune, J.L., Colla, S.R., Coristine, L.E., Davy, C.M., Flockhart, D.T.T., Schuster, R., Orihel, D.M., 2019. Are we accurately estimating the potential role of pollution in the decline of species at risk in Canada? FACETS. https://doi.org/10.1139/facets-2019-0025

McGill, B., 2018. Mining related question.

Mullins, P., Wright, P., 2016. Connecting Outdoor Recreation, Community, and Health in Living Landscapes, in: The Integration Imperative: Cumulative Environmental, Community and Health Impacts of Multiple Natural Resource Developments. Springer International AG.

Newbold, T., Hudson, L.N., Hill, S.L.L., Contu, S., Lysenko, I., Senior, R.A., Börger, L., Bennett, D.J., Choimes, A., Collen, B., Day, J., De Palma, A., Díaz, S., Echeverria-Londoño, S., Edgar, M.J., Feldman, A., Garon, M., Harrison, M.L.K., Alhusseini, T., Ingram, D.J., Itescu, Y., Kattge, J., Kemp, V., Kirkpatrick, L., Kleyer, M., Correia, D.L.P., Martin, C.D., Meiri, S., Novosolov, M., Pan, Y., Phillips, H.R.P., Purves, D.W., Robinson, A., Simpson, J., Tuck, S.L., Weiher, E., White, H.J., Ewers, R.M., Mace, G.M., Scharlemann, J.P.W., Purvis, A., 2015. Global effects of land use on local terrestrial biodiversity. Nature 520, 45–50. https://doi.org/10.1038/nature14324

Nishnawbe Aski Nation, n.d. Canadian Boreal Forest Agreement. Nishnawbe Aski Nation, Thunder Bay, On [WWW Document]. URL http://www.nan.on.ca/article/canadian-boreal-forest-agreement-462.asp (accessed 8.6.20).

NOAA, 2019. Version 1 VIIRS Day/Night Band Nighttime Lights.

O’Donnell, B., 1989. Indian and Non-Native Use of Nitinat Lake and River An Historical Perspective, Native Affairs Division, Policy and Program Planning. Fisheries and Oceans Canada.

Pasher, J., Seed, E., Duffe, J., 2013. Development of boreal ecosystem anthropogenic disturbance layers for Canada based on 2008 to 2010 Landsat imagery. Can. J. Remote Sens. 39, 42–58. https://doi.org/10.5589/m13-007

Prebble, M., Wilmshurst, J.M., 2009. Detecting the initial impact of humans and introduced species on island environments in Remote Oceania using palaeoecology. Biol. Invasions 11, 1529–1556. https://doi.org/10.1007/s10530-008-9405-0

Primack, R.B., 1993. Essentials of Conservation Biology. Sinauer Associates Inc.

Ricketts, T., Imhoff, M., 2003. Biodiversity, Urban Areas, and Agriculture: Locating Priority Ecoregions for Conservation. Conserv. Ecol. 8.

Robb, C.K., 2014. Assessing the Impact of Human Activities on British Columbia’s Estuaries. PLOS ONE 9, e99578. https://doi.org/10.1371/journal.pone.0099578

Sala, O.E., Chapin, F.S., Iii, Armesto, J.J., Berlow, E., Bloomfield, J., Dirzo, R., Huber-Sanwald, E., Huenneke, L.F., Jackson, R.B., Kinzig, A., Leemans, R., Lodge, D.M., Mooney, H.A., Oesterheld, M., Poff, N.L., Sykes, M.T., Walker, B.H., Walker, M., Wall, D.H., 2000. Global Biodiversity Scenarios for the Year 2100. Science 287, 1770–1774. https://doi.org/10.1126/science.287.5459.1770

Sanderson, E.W., Jaiteh, M., Levy, M.A., Redford, K.H., Wannebo, A., Woolmer, G., 2002. The Human Footprint and the Last of the Wild. BioScience 52, 891–904.

Shackelford, N., Standish, R.J., Ripple, W., Starzomski, B.M., 2017. Threats to biodiversity from cumulative human impacts in one of North America’s last wildlife frontiers. Conserv. Biol. 32, 672–684. https://doi.org/10.1111/cobi.13036

Smith, H.C., 1983. Growth of Appalachian Hardwoods Kept Free to Grow from 2 to 12 Years after Clearcutting. Res Pap NE-528 Broomall PA US Dep. Agric. For. Serv. Northeast. For. Experiement Stn. 6p 528.

Steffen, W., Richardson, K., Rockström, J., Cornell, S.E., Fetzer, I., Bennett, E.M., Biggs, R., Carpenter, S.R., Vries, W. de, Wit, C.A. de, Folke, C., Gerten, D., Heinke, J., Mace, G.M., Persson, L.M., Ramanathan, V., Reyers, B., Sörlin, S., 2015. Planetary boundaries: Guiding human development on a changing planet. Science 347, 1259855. https://doi.org/10.1126/science.1259855

Sterling, S.M., Garroway, K., Guan, Y., Ambrose, S.M., Horne, P., Kennedy, G.W., 2014. A new watershed assessment framework for Nova Scotia: A high-level, integrated approach for regions without a dense network of monitoring stations. J. Hydrol. 519, Part C, 2596–2612. https://doi.org/10.1016/j.jhydrol.2014.07.063

Tapia-Armijos, M.F., Homeier, J., Draper Munt, D., 2017. Spatio-temporal analysis of the human footprint in South Ecuador: Influence of human pressure on ecosystems and effectiveness of protected areas. Appl. Geogr. 78, 22–32. https://doi.org/10.1016/j.apgeog.2016.10.007

Tratalos, J., Fuller, R.A., Warren, P.H., Davies, R.G., Gaston, K.J., 2007. Urban form, biodiversity potential and ecosystem services. Landsc. Urban Plan. 83, 308–317. https://doi.org/10.1016/j.landurbplan.2007.05.003

Trombulak, S.C., Frissell, C.A., 2000. Review of Ecological Effects of Roads on Terrestrial and Aquatic Communities. Conserv. Biol. 14, 18–30. https://doi.org/10.1046/j.1523-1739.2000.99084.x

Tulloch, V.J., Tulloch, A.I., Visconti, P., Halpern, B.S., Watson, J.E., Evans, M.C., Auerbach, N.A., Barnes, M., Beger, M., Chadès, I., Giakoumi, S., McDonald-Madden, E., Murray, N.J., Ringma, J., Possingham, H.P., 2015. Why do we map threats? Linking threat mapping with actions to make better conservation decisions. Front. Ecol. Environ. 13, 91–99. https://doi.org/10.1890/140022

van der Marel, R.C., Holroyd, P.C., Duinker, P.N., 2020. Managing human footprint to achieve large-landscape conservation outcomes: Establishing density limits on motorized route-user networks in Alberta’s Eastern Slopes. Glob. Ecol. Conserv. 22, e00901. https://doi.org/10.1016/j.gecco.2019.e00901

Venter, O., Brodeur, N.N., Nemiroff, L., Belland, B., Dolinsek, I.J., Grant, J.W.A., 2006. Threats to Endangered Species in Canada. BioScience 56, 903–910. https://doi.org/10.1641/0006-3568(2006)56[903:TTESIC]2.0.CO;2

Venter, O., Sanderson, E.W., Magrach, A., Allan, J.R., Beher, J., Jones, K.R., Possingham, H.P., Laurance, W.F., Wood, P., Fekete, B.M., Levy, M.A., Watson, J.E.M., 2016a. Sixteen years of change in the global terrestrial human footprint and implications for biodiversity conservation. Nat. Commun. 7, 12558. https://doi.org/10.1038/ncomms12558

Venter, O., Sanderson, E.W., Magrach, A., Allan, J.R., Beher, J., Jones, K.R., Possingham, H.P., Laurance, W.F., Wood, P., Fekete, B.M., Levy, M.A., Watson, J.E.M., 2016b. Global terrestrial Human Footprint maps for 1993 and 2009. Sci. Data 3, 160067. https://doi.org/10.1038/sdata.2016.67

Viera, A.J., Garrett, J.M., 2005. Understanding interobserver agreement: The kappa statistic. Fam. Med. 37, 360–363.

Waller, D., Reo, N., 2018. First stewards: ecological outcomes of forest and wildlife stewardship by indigenous peoples of Wisconsin, USA. Ecol. Soc. https://doi.org/10.5751/ES-09865-230145

Watson, J.E.M., Shanahan, D.F., Di Marco, M., Allan, J., Laurance, W.F., Sanderson, E.W., Mackey, B., Venter, O., 2016. Catastrophic Declines in Wilderness Areas Undermine Global Environment Targets. Curr. Biol. 26, 2929–2934. https://doi.org/10.1016/j.cub.2016.08.049

White, J.C., Wulder, M.A., Hermosilla, T., Coops, N.C., Hobart, G.W., 2017. A nationwide annual characterization of 25 years of forest disturbance and recovery for Canada using Landsat time series. Remote Sens. Environ. 194, 303–321. https://doi.org/10.1016/j.rse.2017.03.035

Williams, B.A., Venter, O., Allan, J.R., Atkinson, S.C., Rehbein, J.A., Ward, M., Di Marco, M., Grantham, H.S., Ervin, J., Goetz, S.J., Hansen, A.J., Jantz, P., Pillay, R., Rodríguez-Buriticá, S., Supples, C., Virnig, A.L.S., Watson, J.E.M., 2020. Change in Terrestrial Human Footprint Drives Continued Loss of Intact Ecosystems. One Earth 3, 371–382. https://doi.org/10.1016/j.oneear.2020.08.009

Woo-Durand, C., Matte, J.-M., Cuddihy, G., McGourdji, C.L., Venter, O., Grant, J.W.A., 2020. Increasing importance of climate change and other threats to at-risk species in Canada. Environ. Rev. er-2020–0032. https://doi.org/10.1139/er-2020-0032

Woolmer, G., Trombulak, S.C., Ray, J.C., Doran, P.J., Anderson, M.G., Baldwin, R.F., Morgan, A., Sanderson, E.W., 2008. Rescaling the Human Footprint: A tool for conservation planning at an ecoregional scale. Landsc. Urban Plan. 87, 42–53. https://doi.org/10.1016/j.landurbplan.2008.04.005

WWF Canada, 2020. Living Planet Report Canada 2020 - Wildlife At Risk. WWF Canada, Toronto, On.

WWF Canada, 2003. The Nature Audit: Setting Canada’s Conservation Agenda for the 21st Century (No. 1). World Wildlife Fund Canada, Toronto, Canada.

