## supplemental files for "Canada’s human footprint reveals large intact areas juxtaposed against areas under immense anthropogenic pressure"

**Supplementary information**

**Supplementary 1**

Visual interpretation of satellite images for mapping human threats (Adapted from Venter et al. 2016a, 2016b)

When interpreting images, interpreters can zoom in, zoom out and pan around to identify threats. For sample areas where only coarse scale Landsat images are available (i.e. 15 m resolution), images can be used if it is deemed that they are sufficient to allow classification for the area, which may be possible in highly green intact areas or in high latitudes where inhabitants are low. Often, when zoomed out in coarse resolution, it is possible to be certain of human threats. If after zooming and panning it is impossible to be certain, the data should be left blank and marked uncertain. For small areas within plots that have identified human threats, these areas are often assigned the land cover category of the wider landscape, i.e. urban, forestry or crops. For example, if there is bare ground across a plot within farmland, it is likely to be tilled farmland, likewise a brown patch in a forested landscape is likely to be a recently felled clear cut. Distinguishing between crops and pasture is a challenge, zooming in to look for linear planting lines or signs of cattle trails or their feeding/drinking points may help. Some land cover types are not mutually exclusive, for instance, urban areas may also be scored as high density for roads and human dwellings. Crops, pasture, urban and forestry are mutually exclusive at a site, but can co-occur within a $1km^2$ or $100km^2$ sample area. Following visual interpretation, interpreters should mark their interpretation as ‘certain’ or ‘not certain’. Certain means that 95% of the time you will be right. The resolution and year of each image will be recorded for all plots, whether or not they have data entered and are certain or not.

The samples are selected using a random sampling. Those are automatically overlaid with ESRI high resolution images within ArcGIS 10.5.1, allowing a rapid access to recent remote sensing images with zooming capabilities. If using ArcMap, ensure the plots are projected on the fly and the basemap is the source determining the projected coordinate system to speed up the process.

**Figure 1** The level of detail of images available in many locations. In the first panel, horses can be seen grazing in front of the farm house, and hay bales can be seen wrapped and stacked to the right of the barn. In the second panel, the uniform grey of concrete, as well as individual containers and the cranes used to move them can be seen. Shape, size, texture and colour are important characteristics for identifying human threats on the environment.

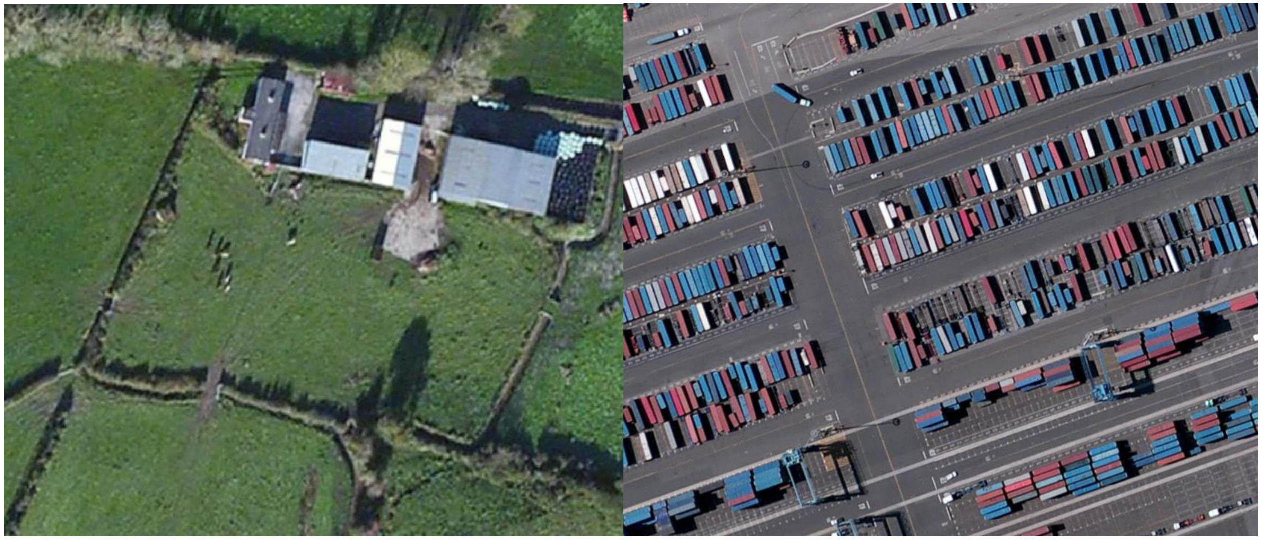

| Pressure | Description | Scoring |
|---|---|---|
| Built Environments | Built environments are human produced areas that provide the setting for human activity. These are primarily urban settings, including buildings, paved land and urban parks, and excludes isolated roads and isolated housing. These also include areas such as airports and unidentified industry. They are easily identified by sharp contrasts in tones, widespread homogeneous grey surfaces, and recognizable human constructed shapes. They are scored based on polygon scoring, a percent of the total image covered. Urban parks, golf courses, shopping centers are all examples of built environments. | None = 0,<br>sparse = 1, <12.5%<br>medium = 2, >12.5%<br>dense = 3, >50% |
| 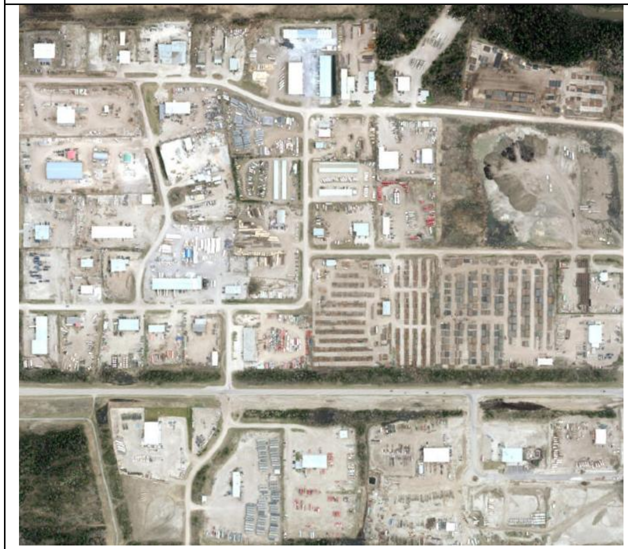 | 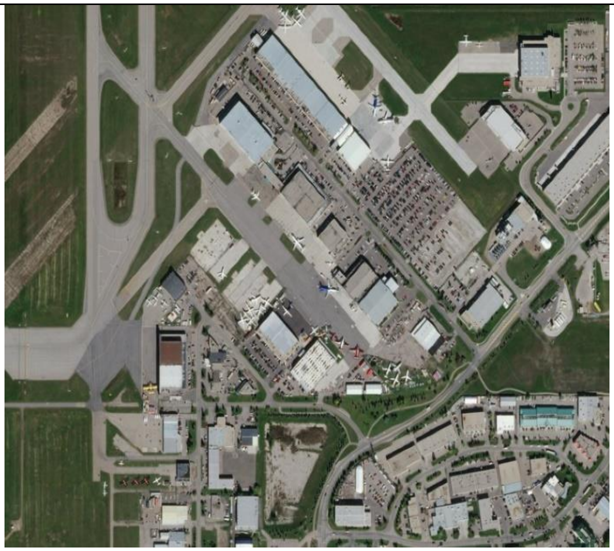 | 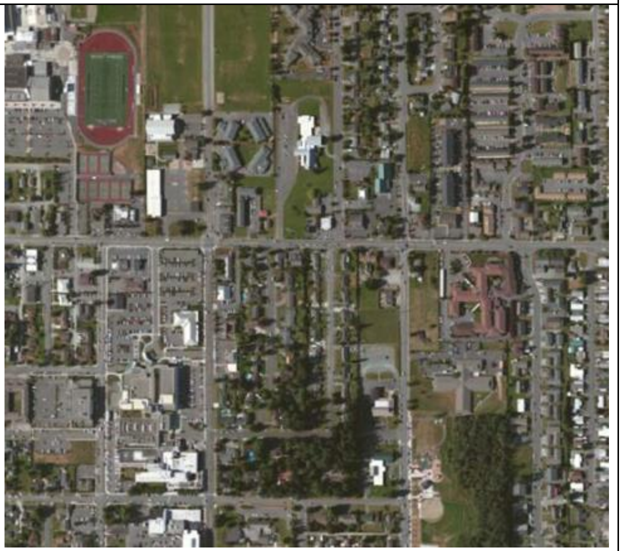 |

| | | |
|---|---|---|
| Crops | Crop lands are cultivated areas used for annual or perennial crops, such as orchards or vineyards. Typically exhibit a checkerboard pattern of crop land pattern from different crop stages (exhibited by varying grey tones) and differences in tillage directions. Crop land areas, generally devoid of trees, possess a smoother texture than pasture land areas and often have linear markings from planting, harvesting or tilling lines. | None = 0,<br>sparse = 1, <12.5%<br>medium = 2, >12.5%<br>dense = 3, >50% |
| 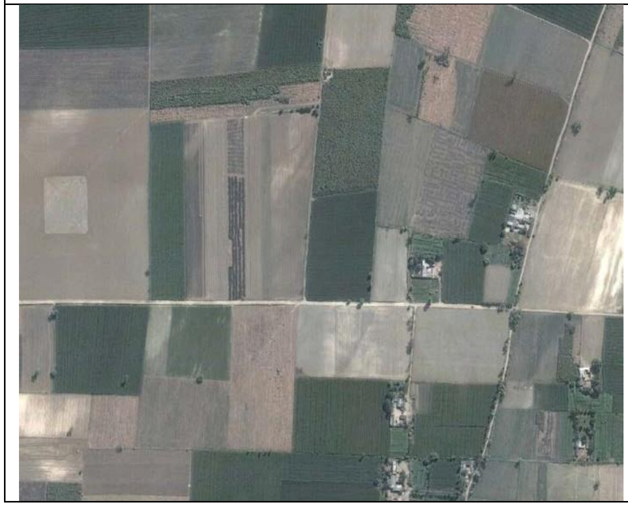 | 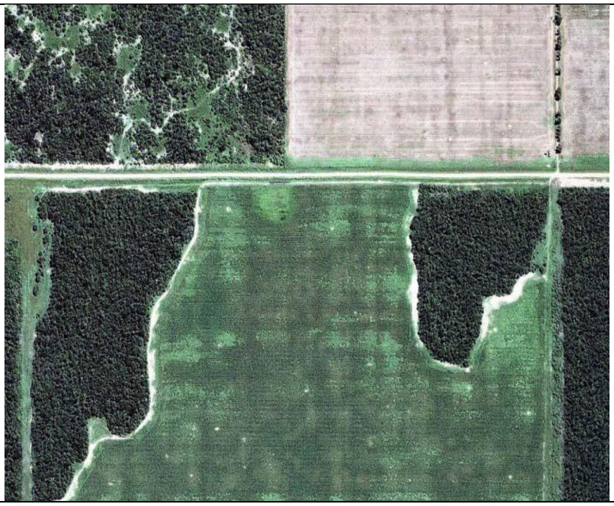 | 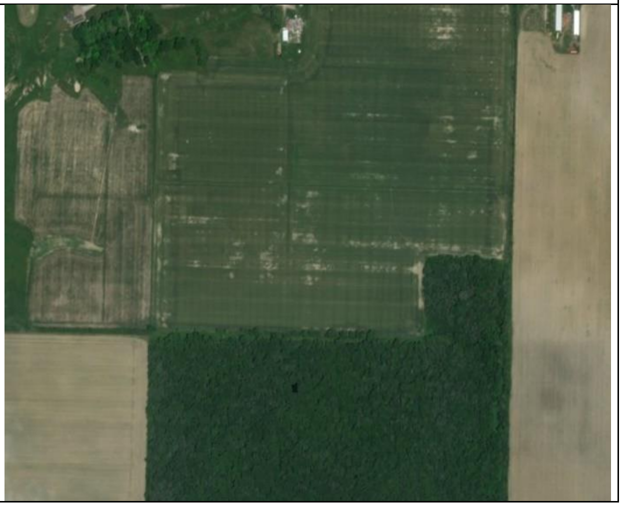 |

| | | |
|---|---|---|
| Pasture | Land covered with grass and grazing animals, especially cattle or sheep. Often characterized by fencing without linear cropping, but often with linear changes in vegetation blocks along fence lines. Cattle or their tracks, as well as vehicle access tracks may be visible. | None = 0,<br>sparse = 1, <12.5%<br>medium = 2, >12.5%<br>dense = 3, >50% |
| 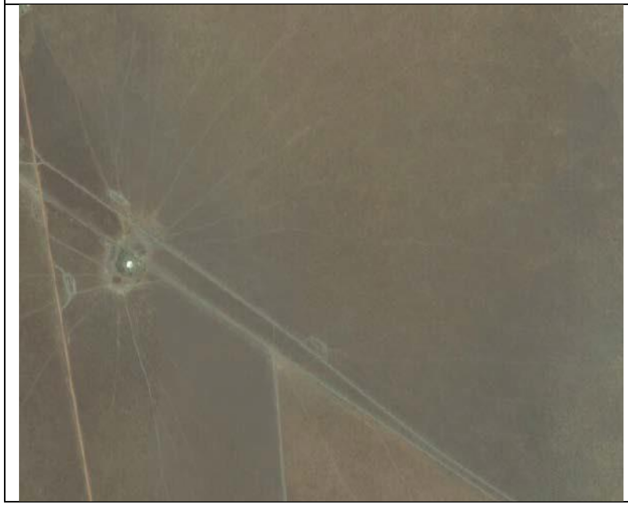 | 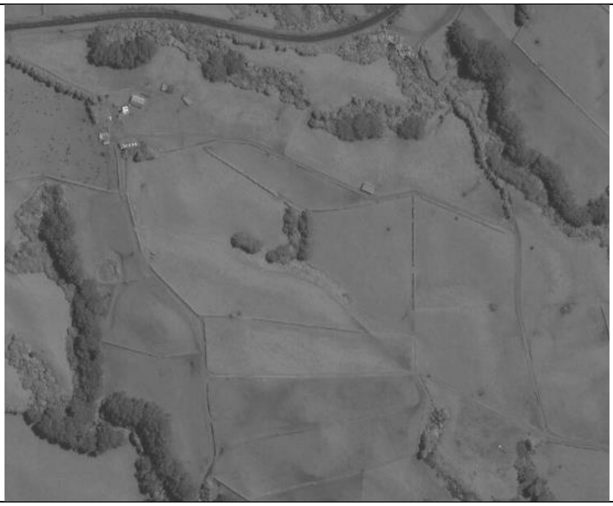 | 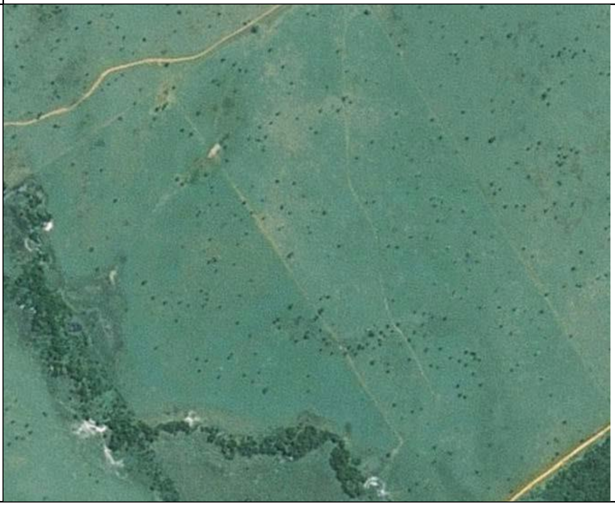 |

| | | |
|---|---|---|
| Roads-paved, unpaved and private | Linear infrastructure with a wide homogeneous grey surface, and often a disturbed vegetation or bare earth band in parallel. Paved roads have a grey surface, unpaved roads have a brown surface. Unpaved logging roads can often appear grey, but can be determined unpaved by proximity to urban centers and general appearance of the surrounding landscape. Private roads are not used for transportation by the public, but rather provide private access, such as access to farm fields. | None = 0,<br>sparse = 1, at least one road visible<br>medium = 2, roads with length that traverses the image twice<br>dense = 3, roads with length that traverses the image 5 times |
| 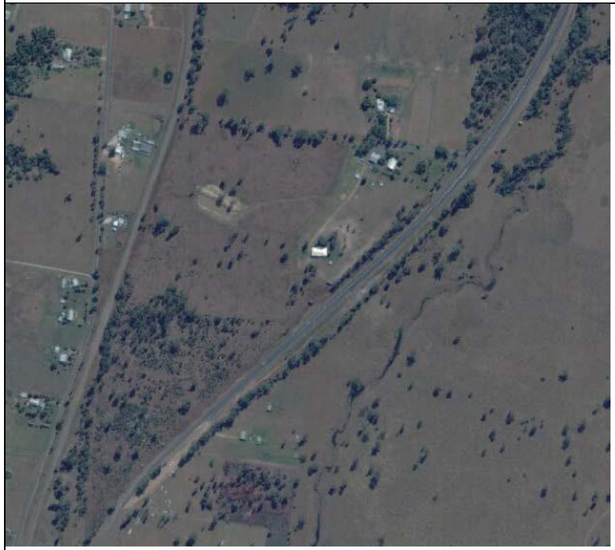 | 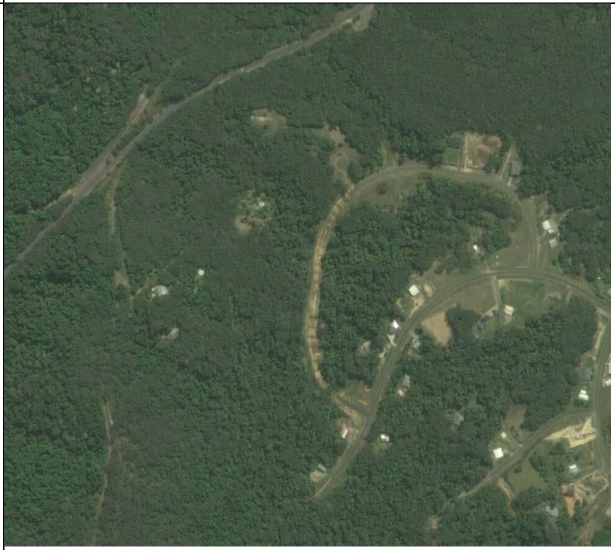 | 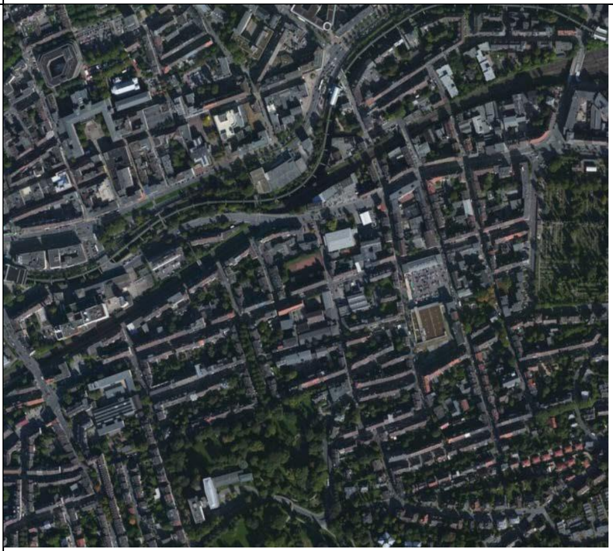 |

| | | |
|---|---|---|
| Forestry | Harvesting of natural or plantation forest. Can be clear- fell harvesting, common in temperate forests, or selective logging. Clear-fell harvesting characterized by large patches of felled forest of often irregular shape following topographic features. Selective harvesting characterized by much smaller harvest patches, a network of dirt roads with noticeable small cleared areas with dirt surface used for landing logging. Plantation forests can be distinguished by their uniform tree cover, and sometimes linear planting rows. If an area has a threat from forestry, it is given an overall score for any and all forestry present there. If there are cutblocks that are recent such that slash piles and bareland is present, or bareland between the young light green trees, this is also given its own score for recent cut forestry. It is possible to have both categories scored as a 3, 2 or 1, or any other combination. | None = 0,<br>sparse = 1, <12.5%<br>medium = 2, >12.5%<br>dense = 3, >50% |
| 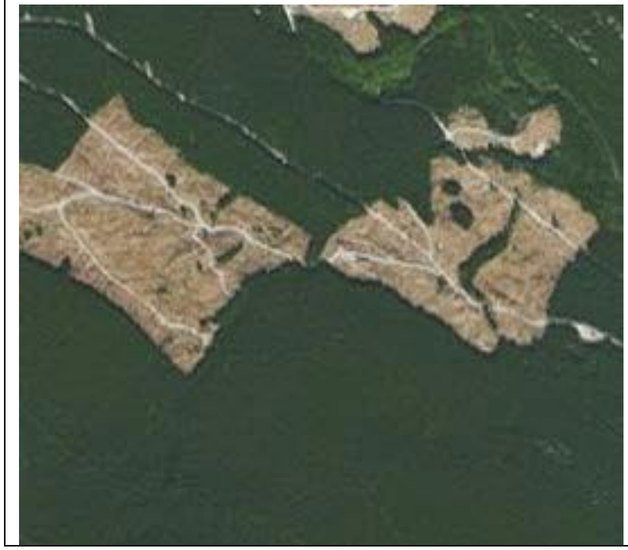 | 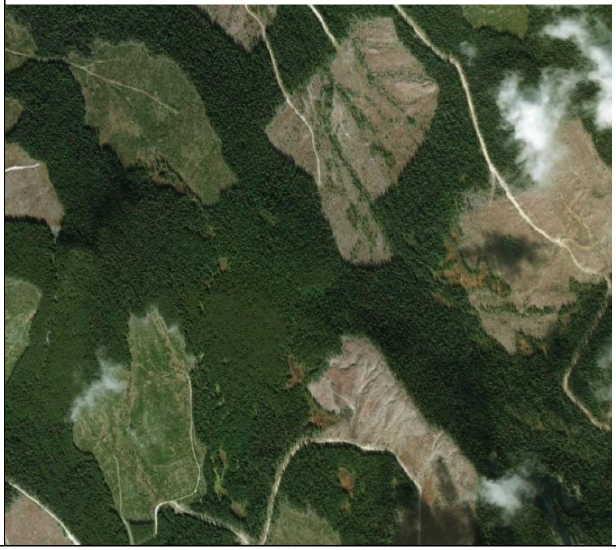 | 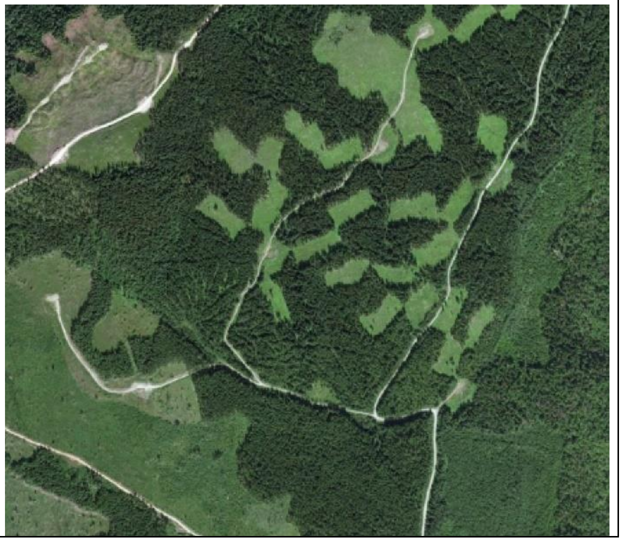 |

| | | |
|---|---|---|
| Human dwellings (looks at population density) | Human dwellings, including dense urban areas with apartment buildings, and sparser suburban and rural housing. It is only being assessed for number of human dwellings and no other infrastructure included. | None = 0,<br>sparse = 1, <4 single-family dwellings per $km^2$ medium = 2, <20 single-family dwellings per $km^2$ dense = 3, >20 dwellings per $km^2$, or 1 apartment building per $km^2$ |
| 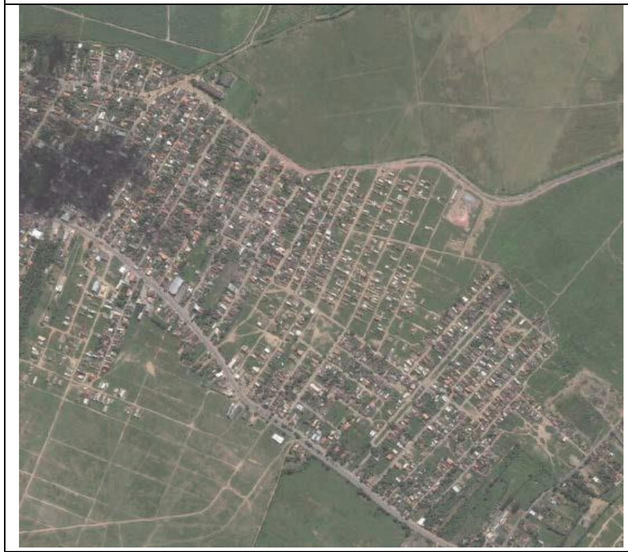 | 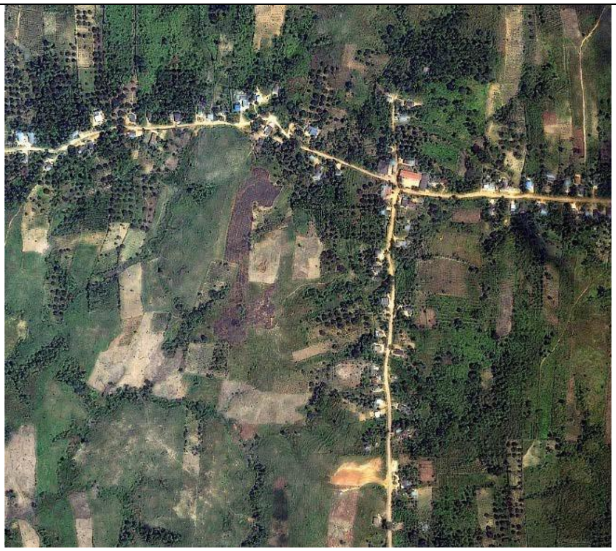 | 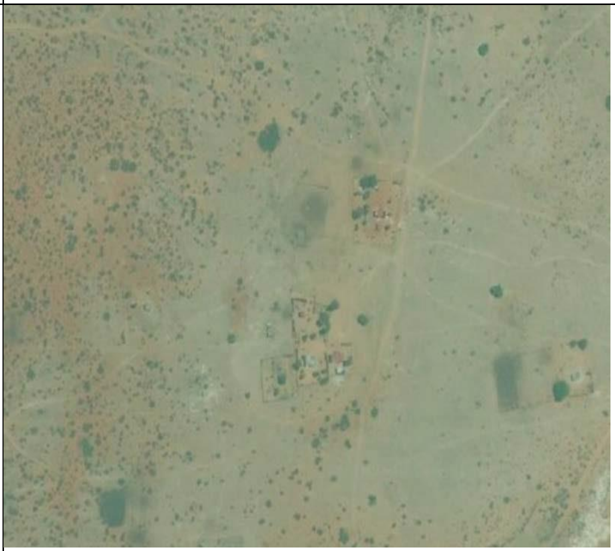 |

| | | |
|---|---|---|
| Navigable waterways | Navigable waterways appear wide and deep enough for a vessel to travel, and lack impassable areas of whitewater. Signs of human activity along the shoreline, such as human structures or roads leading to the water within 40km of the sample plot mean the waterway is likely to be navigated. This threat is scoring for access, not size of water body. If the entire image is water (ie: ocean) it only gets a score of 1 as there is still only one stretch of access points that it provides. | None = 0,<br>sparse = 1, at least one navigable waterway<br>medium = 2, navigable waterways with length that traverses the image twice<br>dense = 3, navigable waterways with length that traverses the image 5 times |
| 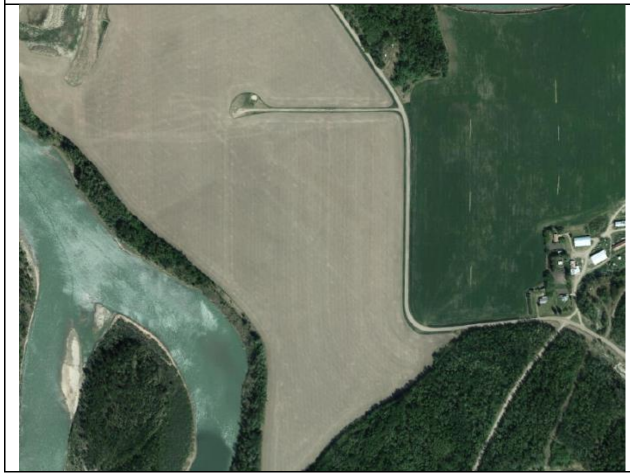 | 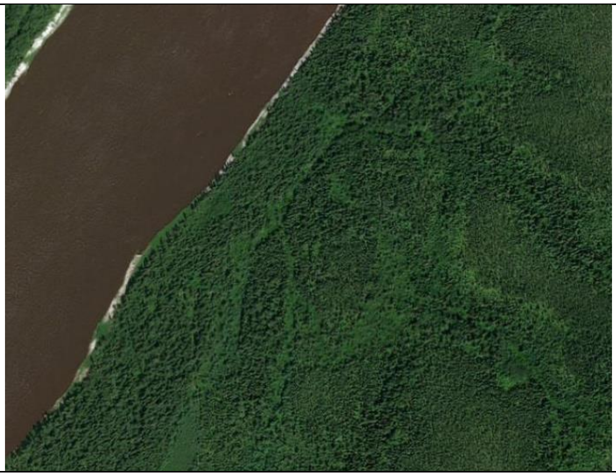 | 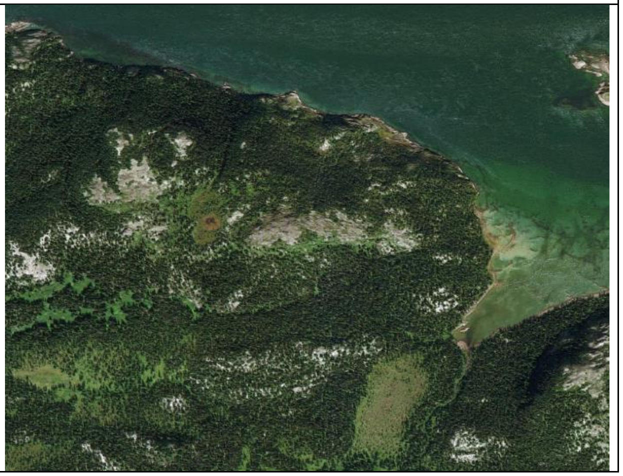 |

| | | |
|---|---|---|
| Railways | Linear infrastructure with a wide homogeneous grey or brown surface, and often a disturbed vegetation or bare earth band in parallel. | None = 0,<br>sparse = 1, at least one railway<br>medium = 2, rail with length that traverses the image twice<br>dense = 3, rail with length that traverses the image 5 times |
| 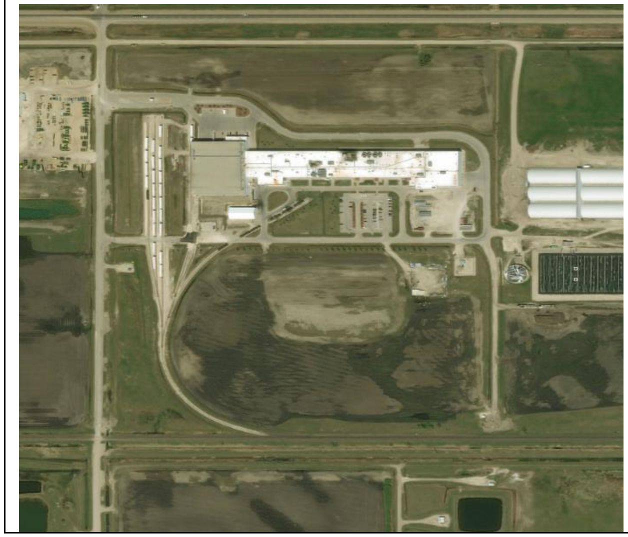 | 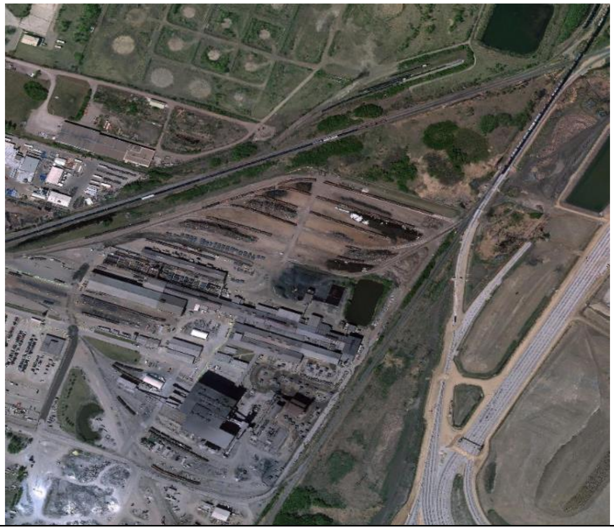 | 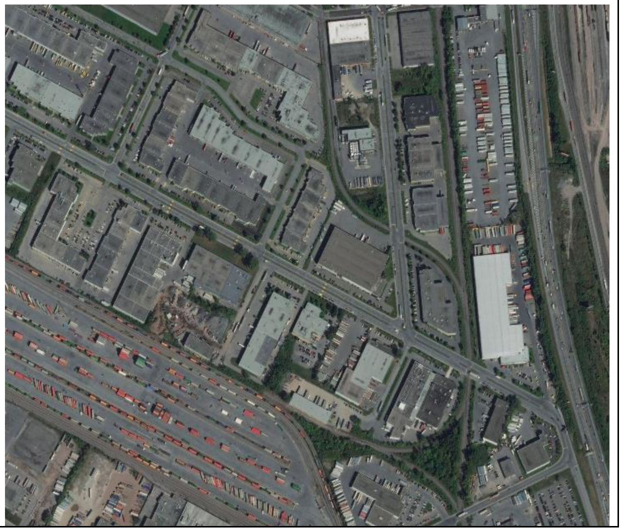 |

| | | |
|---|---|---|
| Dams | Linear infrastructure crossing a body of water, often grey in colour. Often associated with flooding on one side of the dam. | None = 0,<br>sparse = 1, at least one dam visible<br>medium = 2, dams with length that traverses the image twice<br>dense = 3, dams with length that traverses the image 5 times |
| 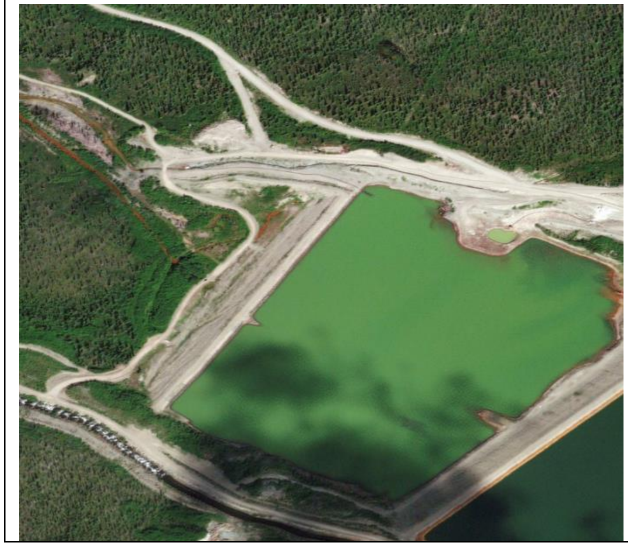 | 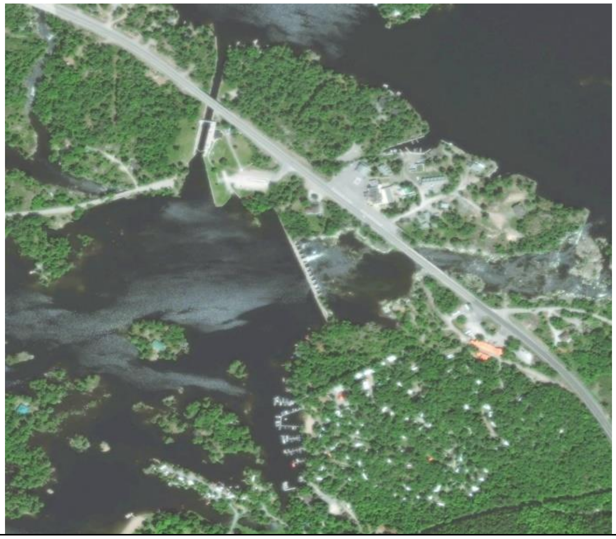 | 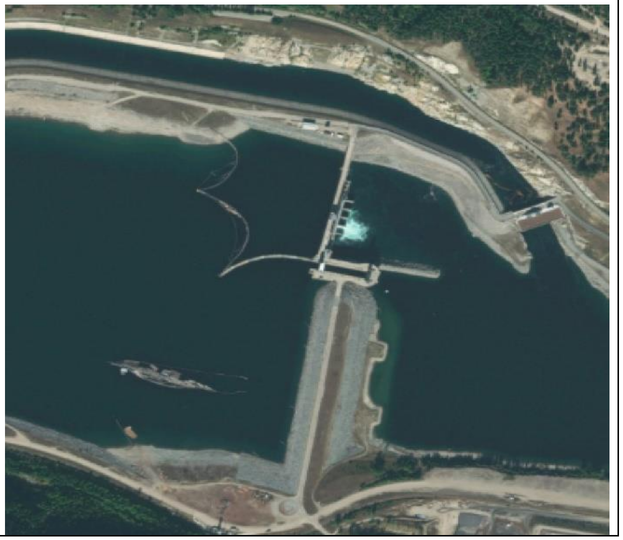 |

| | | |
|---|---|---|
| Mining and associated infrastructure | Areas of mining are often cleared of vegetation and appear grey in colour. Often associated with a few private roads connecting different sites and facilities. | None = 0,<br>sparse = 1, <12.5%<br>medium = 2, >12.5%<br>dense = 3, >50% |
| 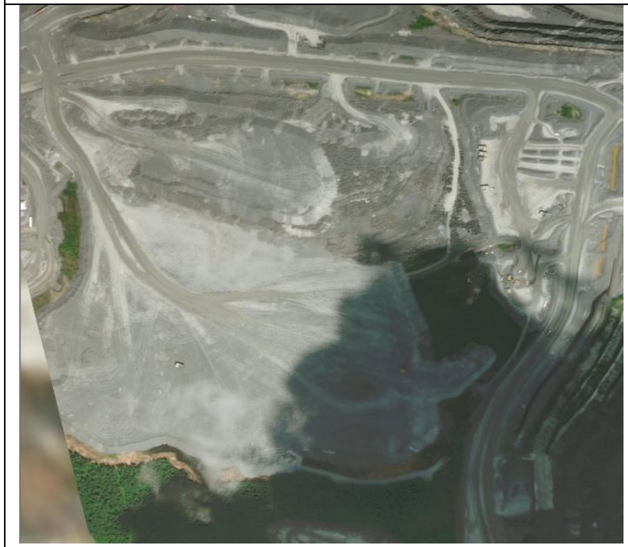 | 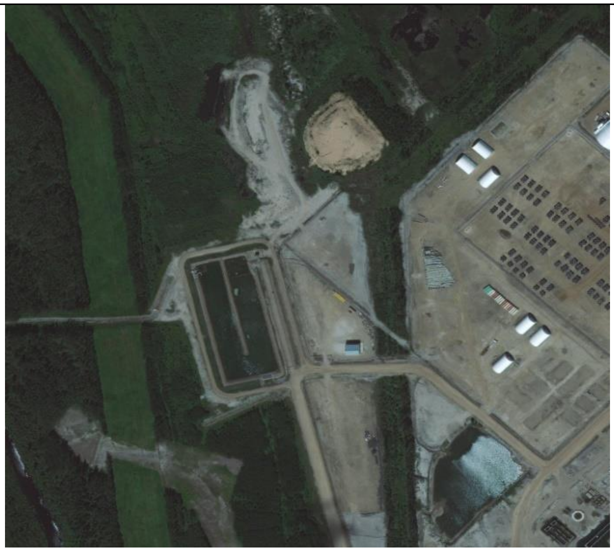 | 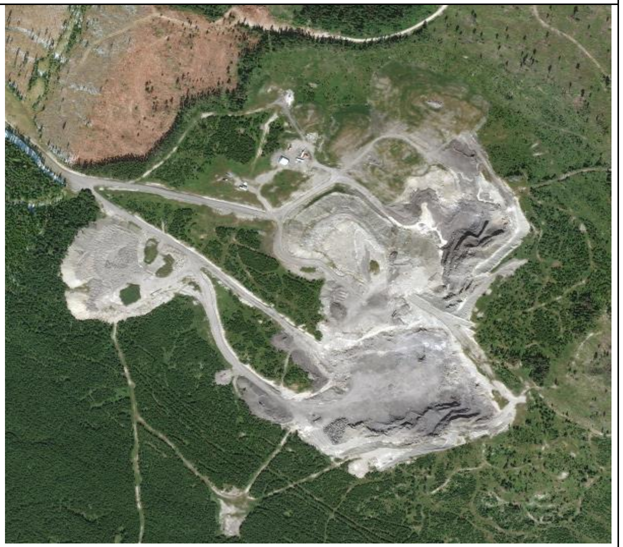 |

| | | |
|---|---|---|
| Oil and gas extraction and associated infrastructure | Series of well pads and connecting linear infrastructure between areas. Often light grey as vegetation has been cleared from surrounding area. Two columns exist for this: linear infrastructure and polygon infrastructure (well pads, buildings, parking lots etc.) and should be scored separately. | Linear: None = 0,<br>sparse = 1, at least one linear oil and gas feature<br>medium = 2, linear oil and gas feature with length that traverses the image twice<br>dense = 3, linear oil and gas feature with length that traverses the image 5 times<br><br>Polygon: None = 0,<br>sparse = 1, <12.5%<br>medium = 2, >12.5%<br>dense = 3, >50% |
| 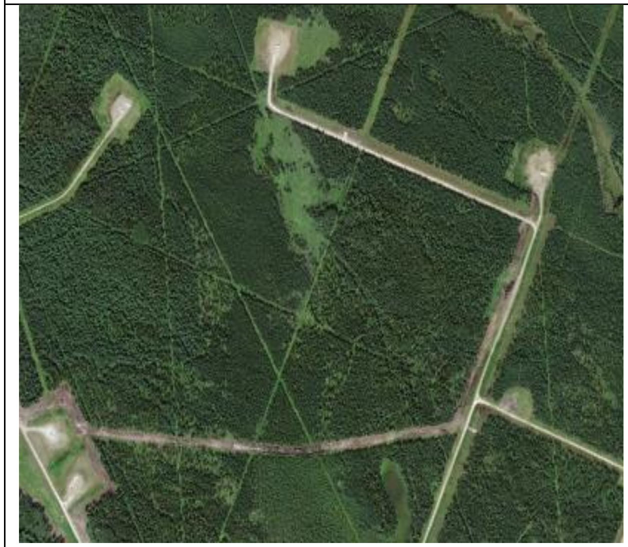 | 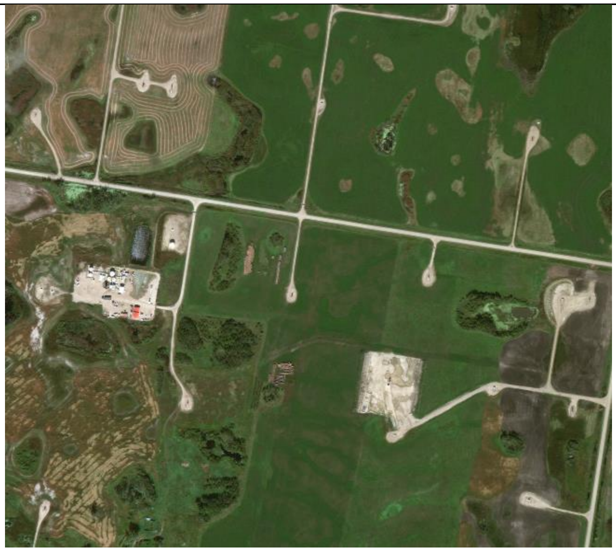 | 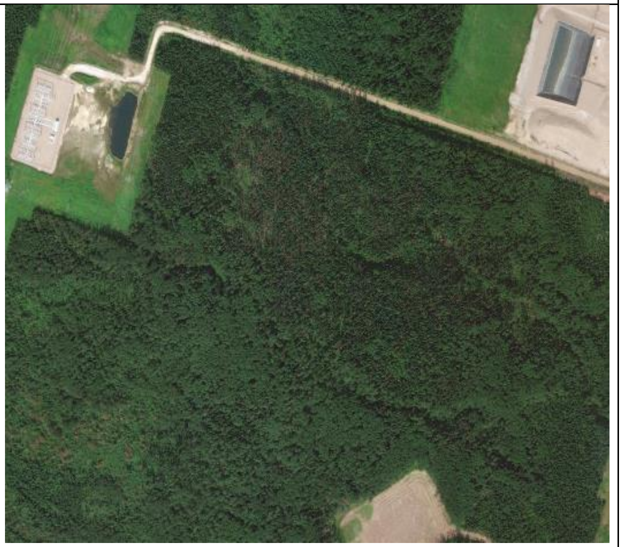 |

| | | |
|---|---|---|
| Seismic | Linear features of cleared land. Often narrow, perfectly straight and numerous. If followed for several kilometres, they may end abruptly. Associated with oil and gas but not always. Can be found in any landscape. Can run right through features such as lakes, but also can be wavy and seen to go around important ecological features. Minimum width is 1.5 metres and maximum is typically 8 metres for older seismic lines. | None = 0,<br>sparse = 1, at least one seismic line visible<br>medium = 2, seismic lines with length that traverses the image twice<br>dense = 3, seismic line with length that traverses the image 5 times |
| 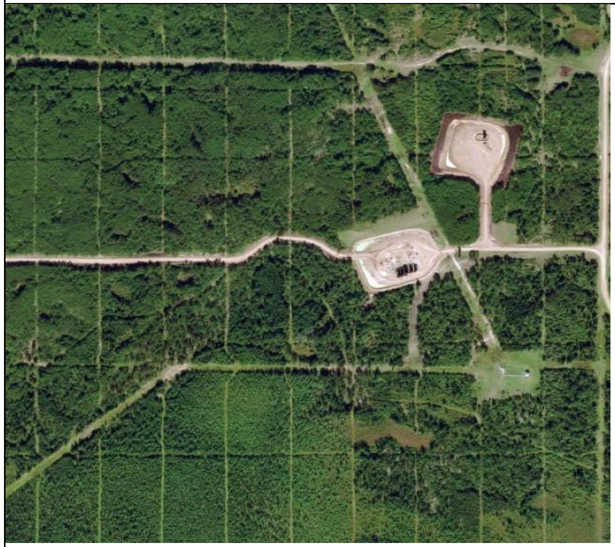 | 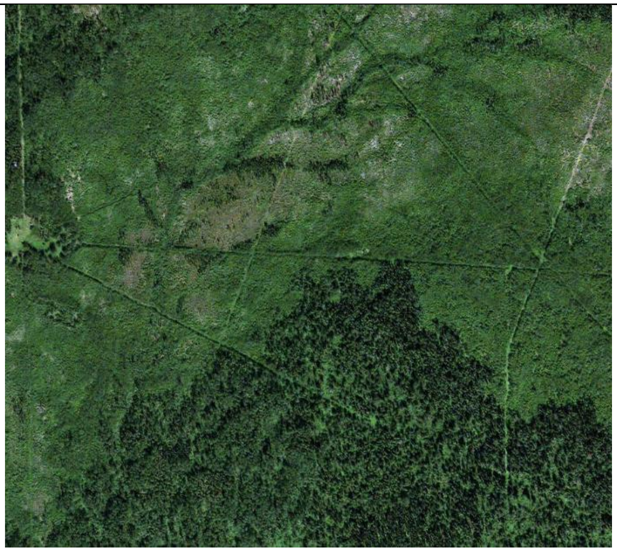 | 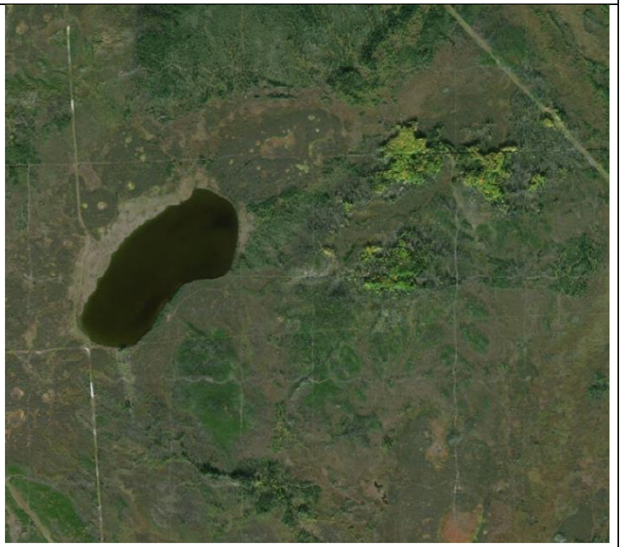 |

| | | |
|---|---|---|
| Electrical infrastructure | Linear swath of cleared vegetation to support the passing of a power line or electrical infrastructure. | None = 0,<br>sparse = 1, at least one transmission line or utility feature visible<br>medium = 2, transmission line or utility feature with length that traverses the image twice<br>dense = 3, transmission line or utility feature with length that traverses the image 5 times |
| 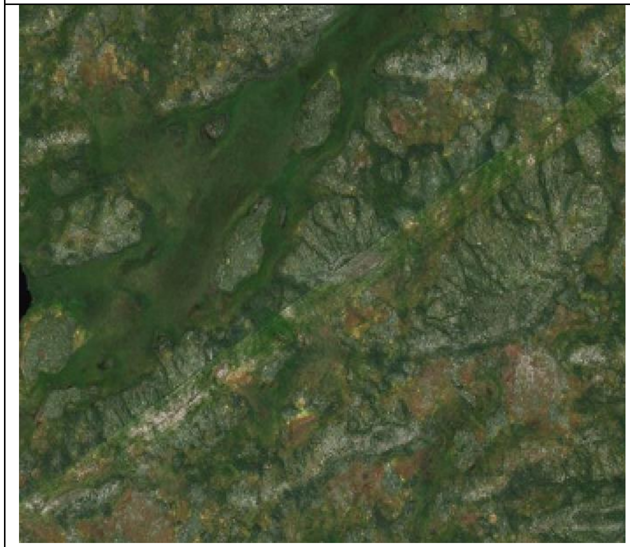 | 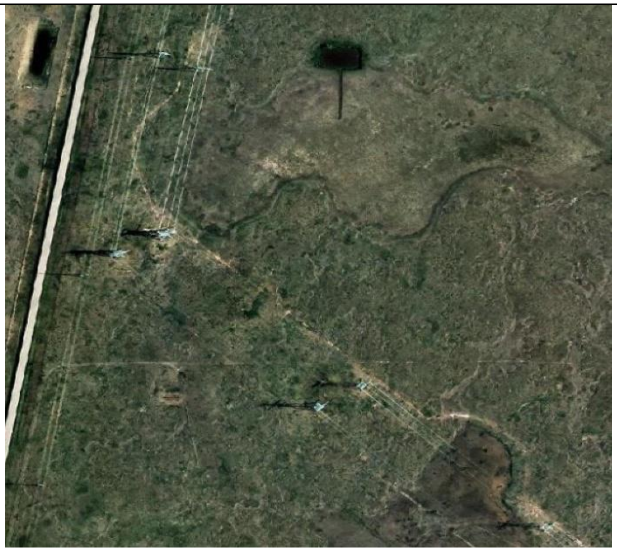 | 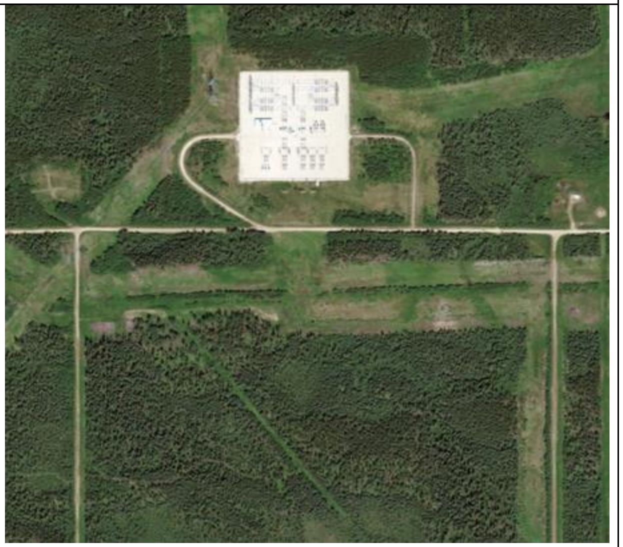 |

**Supplementary 2**

Comparison between the Canadian human footprint (a) and the Global human footprint clipped to Canada (b).

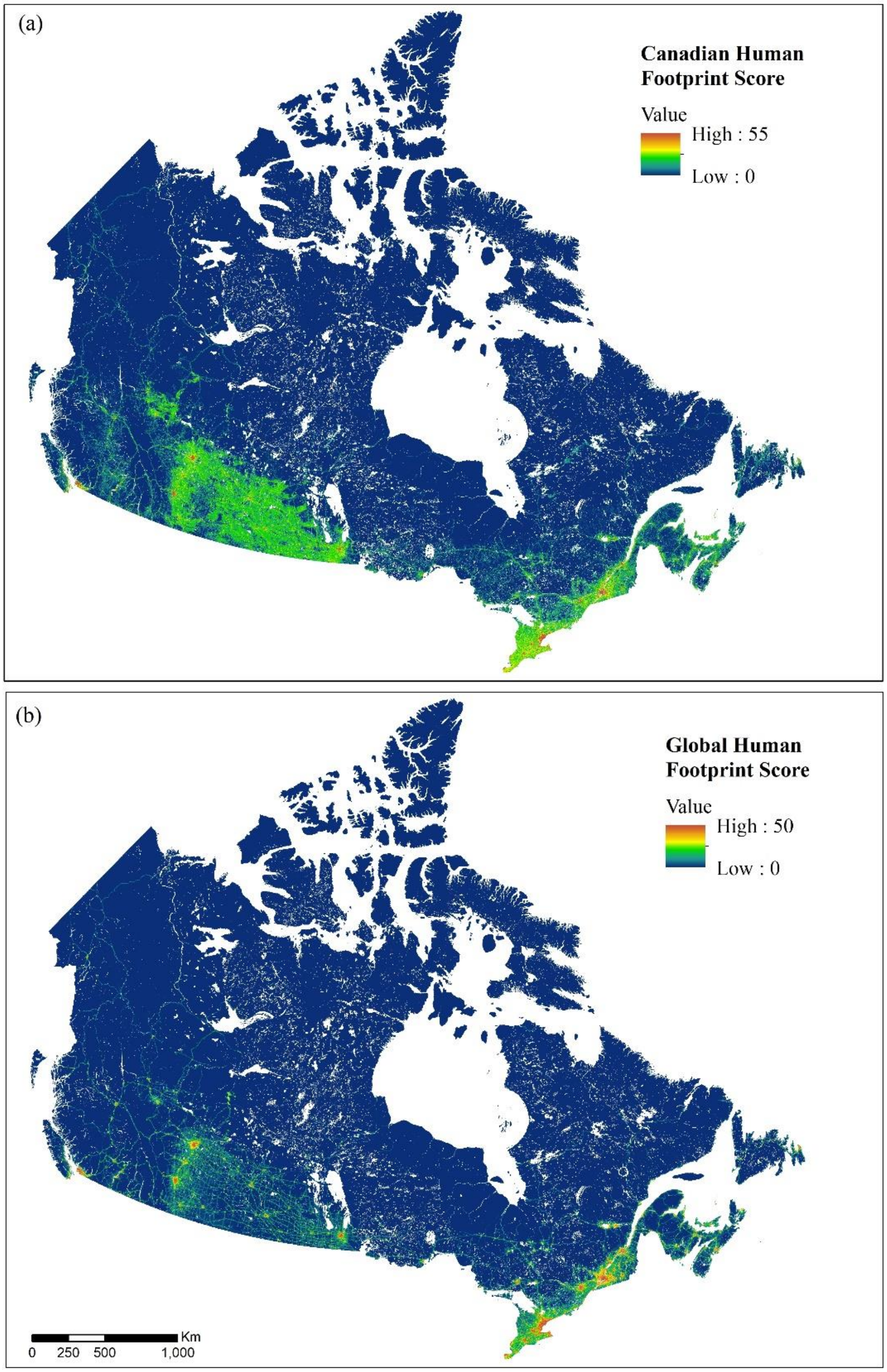

**Supplementary 3**

Sensitivity analysis of the Cohen Kappa Statistic (y-axis) with the Percent Agreement (x-axis) between visual score and Canadian human footprint value. The strength of agreement ranges, appearing as the coloured dashed lines, are as defined by Landis & Koch (1977).

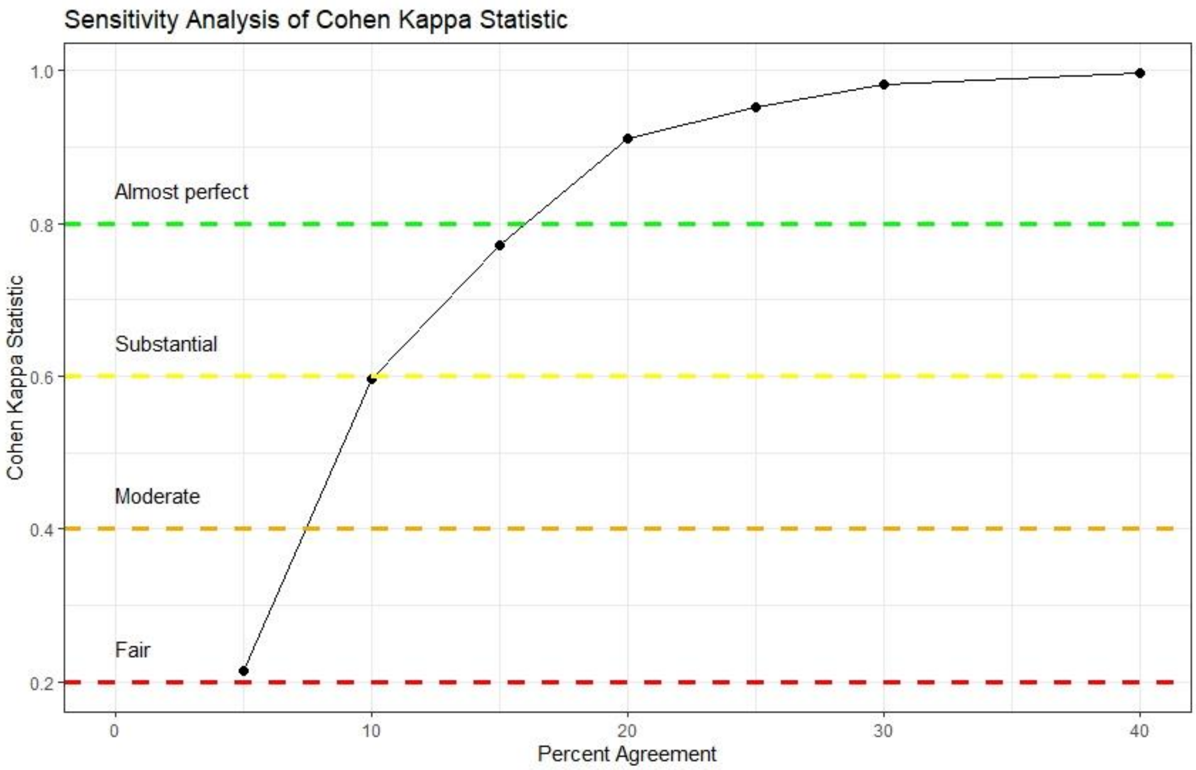
